# Comparative Evaluation of Bioinformatic Pipelines for Full-Length Viral Genome Assembly

**DOI:** 10.1101/2024.03.13.584779

**Authors:** Levente Zsichla, Marius Zeeb, Dávid Fazekas, Éva Áy, Dalma Müller, Karin J. Metzner, Roger Kouyos, Viktor Müller

## Abstract

The increasingly widespread application of next-generation sequencing (NGS) in clinical diagnostics and epidemiological research has generated a demand for robust, fast, automated, and user-friendly bioinformatic workflows. To guide the choice of tools for the assembly of full-length viral genomes from NGS datasets, we assessed the performance and applicability of four widely adopted bioinformatic pipelines (shiver - for which we created a user-friendly Dockerized version, referred to as dshiver; SmaltAlign, viral-ngs, and V-pipe) using both simulated datasets and real-world HIV-1 paired-end short-read sequences and default settings.

All four pipelines produced high-quality consensus genome assemblies and minority variant calls when the reference sequence used for assembly had high similarity to the analyzed sample. However, while shiver and SmaltAlign showed robust performance also with more divergent samples (non-matching subtypes), viral-ngs and V-Pipe proved to be sensitive to genetic distance from the reference sequence. With empirical datasets, SmaltAlign and viral-ngs exhibited substantially shorter runtime compared to V-Pipe and shiver. In terms of applicability, V-Pipe provides the broadest functionalities; SmaltAlign and dshiver combine user-friendliness with robustness; while the use of viral-ngs requires a less computational resources compared to other tools.

To conclude, all four pipelines can perform well in terms of quality metrics; however, the reference sequence needs to be adjusted to closely match the sample data for viral-ngs and V-Pipe. Differences in user-friendliness and runtime may guide the choice of the pipeline in a particular setting. The new Dockerized version of shiver offers ease of use in addition to the accuracy and robustness of the original pipeline.

## Introduction

The clinical diagnostics and molecular epidemiology of viral infections rely increasingly on next-generation sequencing (NGS) due to its speed, high-throughput performance and cost-effectiveness. The Advanced Molecular Detection program in the US applies NGS technologies in nearly every area of infectious disease public health [1], a largely NGS based surveillance of the Covid-19 pandemic yielded more than 8 million full-length SARS-CoV-2 genomic sequences [2], and HIV-1 genotypic resistance testing is also transitioning to NGS technologies in clinical diagnostics [3–5]. NGS has also simplified the sequencing of complete viral genomes, allowing for the detection of mutations outside the traditional target regions for drug resistance genotyping [6–12], and for co-receptor tropism prediction [13–17]. The use of full-length genomes also facilitates the identification of transmission clusters [18,19], subtyping [20], and the detection of recombinant forms [21,22].

Notably, consensus genomes assembled from NGS data can be used as input data for all tools developed for Sanger sequences [23], while minority variant calling (from reads mapped to the consensus sequence) enhances drug resistance prediction [24–27], and enables more precise assessments of infection recency [28] or transmission patterns [29–31].

Genome assemblers typically rely on two traditional approaches: reference-based and *de novo* genome assembly. Reference-based assembly involves mapping sequencing reads to an existing reference sequence and identifying the majority base at each genomic position. This approach is faster, computationally more efficient, and robustly generates full-length genomic sequences by imputation where coverage is low. However, this method is sensitive to the choice of reference and may introduce a bias into read mapping by discarding reads that are divergent from the reference sequence, potentially missing novel information in the sample. In contrast, by constructing the consensus sequence autonomously from the overlapping reads, *de novo* assembly avoids such bias, but is unable to bridge gaps if non-overlapping contigs are produced.

Recently, several bioinformatic pipelines have been developed to combine the advantages of both methods [32–35], but these genome assembly tools have not been comprehensively evaluated. In this work, we assessed the performance of four widely adopted bioinformatic pipelines (shiver, SmaltAlign, viral-ngs, and V-pipe) used for assembling full-length viral genomes from Illumina short reads. In addition to measures of genome quality, we also evaluated technical benchmarks that affect practical utility (runtime and memory usage). To ensure a comprehensive evaluation, we utilized both simulated and empirical HIV-1 datasets.

## Methods

### Workflow

#### Construction of benchmarking references

For each simulated or empirical sample we constructed the consensus sequence of all variants present in the original sample to use as a benchmarking reference (Figure 1). We performed multiple alignment on the simulated SIM and the empirical SGS-FULL sequence sets (see Datasets) using MAFFT [36], called consensus using the *cons* method of the EMBOSS package [37], then cropped the LTR regions, yielding 8500-8700bp long near-full-length HIV-1 genomes. For the analysis of the SS+NGS dataset, we used the Sanger sequences as benchmarking reference.

**Figure 1:**
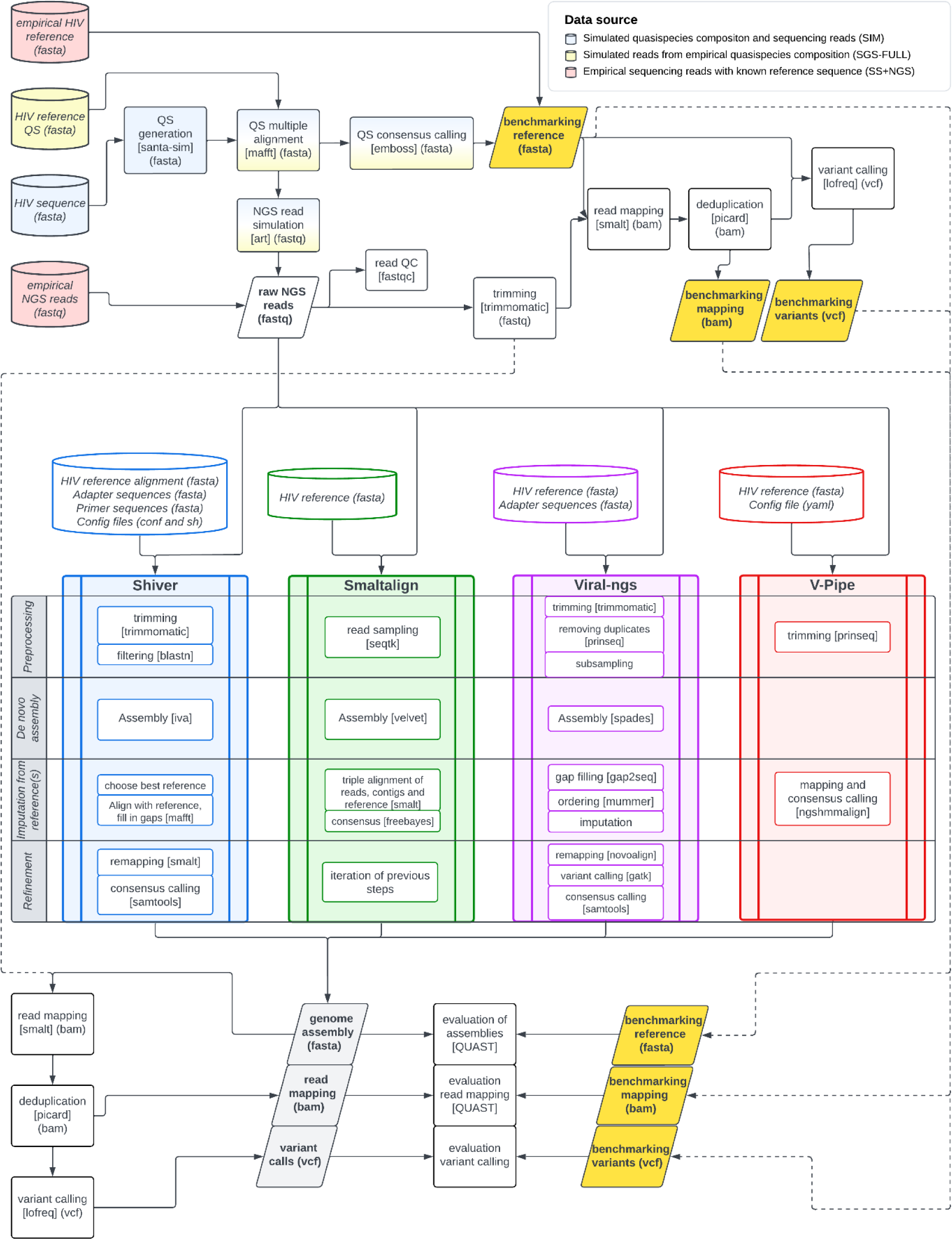
Detailed workflow of the benchmarking pipeline. Data sources are depicted as cylindrical boxes, analysis steps as rectangles, and important files as parallelograms. Applied tools are indicated within square brackets, and file extensions are indicated in parentheses. Differences in the preprocessing of the employed datasets are highlighted with red (SS+NGS), yellow (SGS-FULL), and blue (SIM) coloring. Benchmarking references are emphasized in gold, and assembly results are shown in gray. Abbreviations: QS – quasispecies, QC – quality control

From the SIM and SGS-FULL sequence sets we generated *in silico* Illumina Miseq paired-end short reads with varying fold coverage (500 and 10000 reads/base for the SIM and 2000 reads/base for the SGS-FULL datasets), a read length of 250bp, with Phred quality scores (Q) set between 20 and 40, and a mean fragment size of 700bp using the NGS read simulator ART [38].

We employed Trimmomatic [39] to remove low-quality regions (Q<20) [40] at the beginning and end of the reads (using a sliding window of 4 bases). Reads shorter than 50bp were discarded. We used SMALT to map trimmed reads to the benchmarking reference [41] with an exhaustive search for read mapping, and Picard’s *MarkDuplicates* method [42] to remove any duplicate reads. We inferred single nucleotide polymorphisms and indels from the reference-mapped processed reads using LoFreq [43].

#### Genome assemblers

We evaluated the performance of four established viral genome assembly pipelines: shiver [32] (v1.4.3, for which we created a ‘dockerized’ version to enable easy use: dshiver version v1.4.3_1.0), SmaltAlign (v1.1.0) [34], viral-ngs (v1.25.0) [35,44], and V-pipe (v2.99.3) [45]. The first three pipelines are state-of-the-art tools in viral genome assembly, using a combination of *de novo* and reference-based assembly. They either rely on a reference-based ordering of contigs and subsequent imputation of missing regions (shiver and viral-ngs) or on the iterative mapping of reads to the contigs and a reference sequence (SmaltAlign). The fourth pipeline, V-Pipe, uses a high-precision read mapper specifically designed for viral NGS data (ngshmmalign) to map the reads to a user-specified reference sequence. Figure 1 and Table 1 provide a comparative overview of the analyzed pipelines. Parameters were set to default except for the SS+NGS dataset, where certain parameters in the shiver and viral-ngs pipelines had to be adjusted due to short sequence lengths.

**Table 1:** Comparison of the functionalities of the examined pipelines. Parallel computing is examined at both the pipeline (usage of multiple cores) and batch analysis (parallel computing of multiple samples) levels. Abbreviations: EM – environment management, V – version, DN – de novo assembly, RB – reference-based assembly, RC – reference choice, QP – quality preprocessing, CF – contamination filtering, PC – parallel computing

| NAME | EM | V | DN | RB | RC | QP | CF | PC |
| --- | --- | --- | --- | --- | --- | --- | --- | --- |
| SHIVER | - / VirtualBox / Docker<sup>1</sup> | v1.4.3 | + | + | + | + | + | -/- |
| SMALTALIGN | Conda | v1.1.0 | + | + | - | - | - | +/- |
| VIRAL-NGS | DNAexus / Conda / Docker / Snakemake | v1.25.0 | + | + | - | + | - | +/+ |
| V-PIPE | Snakemake (Conda) | v2.99.3 | - | + | - | + | - | +/+ |

<sup>1</sup> The original version of shiver can be installed without an environment management system using an installation script and is also available as a Virtualbox image. We have created a containerized version (dshiver) described in this paper.

We applied the same steps of genome cropping, read mapping, deduplication, and variant calling to the output consensus genomes.

#### Dshiver: a containerized version of the shiver pipeline

To enable effortless use of the shiver pipeline [32] (also by non-bioinformaticians) we packed it into a Docker container [46] that allows simple installation (on both Linux-based and Windows operating systems) and convenient access to the major capabilities of shiver, supplemented with an improved command line interface, and detailed documentation aimed at readers with basic computer skills for easy use of the pipeline. Furthermore, we eliminated the Python 2 dependency of shiver by updating all scripts to Python 3, to facilitate continuing support and easier integration of shiver into larger pipelines. We also added an automated drug resistance report based on the analysis of the consensus sequence using the Stanford HIVdb algorithm [47]. We call this modified version dshiver (Dockerized version of shiver, v1.4.3_1.0).

The code and the manual of dshiver are publicly available at https://github.com/hcovlab/dshiver and a ready-to-use docker image can be downloaded at https://github.com/orgs/hcovlab/packages/container/package/dshiver.

#### Evaluation of system requirements and the quality of assembled genomes

We assessed the performance of each pipeline using QUAST [48], a dedicated tool for the comparison and quality assessment of genome assemblies. We selected metrics that encompass measurements of genome completeness (fraction of genome assembled) and several aspects of genome accuracy, like mismatch (SNP) and indel rates, the fraction of unidentified bases and the number of misassemblies. Local and global misassemblies were identified using breakpoints, where the left and right flanking sequences overlapped or were separated by a gap (between 200 bp and 1 kbp for local and greater than 1 kbp for global) or were positioned on a different strand. Furthermore, we evaluated mapping precision (difference in the rate of mapped and properly paired reads compared to the benchmark) and the accuracy of minority variant calls using precision (ratio of true positives out of all positive predictions), recall (ratio of true positives correctly identified as positives) and F1 scores (harmonic mean of precision and recall).

To compare the runtime and maximum memory usage of each genome assembler, we performed all analyses on the same computer, with a configuration consisting of an Intel(R) Core(TM) i7-7700 CPU @ 3.60GHz processor, 16GB of system memory and a Ubuntu 22.04.2 LTS operating system. The measurements of CPU time, CPU usage and maximum resident set size (maximum physical memory used) were obtained with the GNU ‘time’ command.

All computer code is publicly available at https://github.com/hcovlab/ViralNGSBenchmarking.

### Datasets

#### Simulated HIV-1 quasispecies composition (SIM)

To generate an *in silico* population of full-length HIV-1 genomic sequences, we utilized SANTA-SIM [49] with initial user-supplied sequences described below. Our customized HIV-1 configuration implemented point mutations (mean 2.5x10^-5^/base/generation [50] multiplied by relative *in vivo* mutation rates for all 12 nucleotide combinations [51]), indels (3x10^-5^/base/generation [52]) and recombination events (0.1 dual infection and 1x10^-5^ recombination per generation [53]) with a constant effective population size of 1000 [54]. The simulations spanned a randomly selected duration between 50 and 1500 generations to generate samples exhibiting diversity patterns resembling both recent and chronic infections.

To cover the genetic variability of HIV-1, we selected four distinct HIV-1 group M subtype consensus sequences (A1, B, C, and CRF01_AE) and one group O consensus sequence (from the HIV-1 consensus sequence alignment of the LANL HIV database [55,56]) for *in silico* quasispecies simulation. This selection also enabled us to examine the effect of an increased divergence of the analyzed sample from the reference genome on the performance of the pipelines (some of which use the HXB2, GenBank: K03455.1, isolate as a fixed reference sequence). Additionally, to show whether providing a well-matched reference sequence alleviates the effect, we included an additional scenario in which genome assemblers used the consensus group O sequence as a reference for the assembly of the *in silico* group O samples. To incorporate further crucial aspects of laboratory work, we also added two different read coverage scenarios (500 and 10,000 per base coverage), and the absence or presence of laboratory contamination into our simulations. The presence of laboratory contamination was simulated by introducing 8,600 randomly generated read pairs, which represented approximately 5% and 100% of the number of reads in the high and low coverage scenarios, respectively. These read pairs were derived from a 55kb fragment of human chromosome 19 (GRCh38.p14, chr22:58283717-58338638) including the 11kb Endogenous Retrovirus Group K3 Member 1 (*ERVK-1*) gene. This procedure resulted in 24 unique parameter combinations (5+1 subtype/reference, 2 coverage and 2 contamination scenarios) with 20 replicates each, totaling 480 simulations.

#### Single genome sequencing data (SGS-FULL)

Single genome amplification and sequencing (SGS) is used to characterize the within-host genetic diversity of chronic HIV infections [57]. The selected SGS dataset contained 13-55 intact near-full-length (∼8800bp) subtype B genomic sequences from each of 5 patients on antiretroviral therapy, four of whom were undergoing treatment interruption during the study period [58].

#### Sanger and next-generation sequencing from the same sample (SS+NGS)

The results of Sanger sequencing, due to the low per base error rate of the instrument (0.0001%) [59], can serve as benchmarks for assessing genome assemblies generated from short-read NGS data. We used unpublished sequence data obtained from parallel Sanger and next-generation sequencing of the same samples to enable such comparisons.

Plasma samples were collected from ART-naive patients diagnosed as HIV-positive between 2016 and 2022 at the Center for HIV, Central Hospital of Southern Pest, National Institute of Hematology and Infectious Disease, Budapest, Hungary. HIV-1 RNA extraction, the amplification of protease, reverse transcriptase and integrase regions and details of the Sanger sequencing method were described previously [60–62]. Additionally, 41 samples were selected for next-generation sequencing using a protocol developed for amplification of near-full-length HIV-1 genome and short-read sequencing based on previous publications [40] (see Supplementary Materials), resulting in 46 NGS datasets from 41 samples. The subtype distribution of sequences determined by REGA (v3.46) [63] from the consensus sequences produced by dshiver was as follows: B (22.0%), CRF 01_AE (9.76%), F1 (7.3%), CRF 19_cpx (4.9%), B-like (4.9%), A1 (2.4%), C-like (2.4%) and other recombinant forms (46.3%).

In sequence positions where Sanger sequencing produced ambiguous results or NGS data strongly indicated a different base call (allele frequency over 0.7), we corrected the sequence to prevent bias in benchmarking results due to inconsistencies between the two datasets.

#### Next generation sequencing dataset for runtime benchmarking (NGS-FULL)

We utilized a publicly available NGS dataset [40] to investigate how the computational demands of each genome assembler scale with increasing dataset size. The chosen dataset comprises NGS data from 92 plasma samples, each subjected to amplification in four overlapping segments covering nearly the entire HIV-1 genome (∼8800bp). Subsequently, the samples were sequenced using the Illumina MiSeq platform, generating 175,302 to 1,649,546 paired-end reads (2x250bp) per sample.

## Results

### Comparison of in vivo and in silico quasispecies datasets

We assessed the performance of four genome assemblers – shiver, SmaltAlign, viral-ngs, and V-Pipe – focusing on assembly quality, precision of read mapping and minority variant calling, and computational resource usage across three distinct HIV-1 datasets. The first encompassed 480 *in silico* HIV-1 quasispecies sequence sets and sequencing reads, spanning 5 viral (sub)types and exploring two coverage and two contamination scenarios (SIM). The second dataset comprised 5 samples from 5 patients, consisting of full-length SGS results combined with simulated sequencing reads (SGS-FULL). The third dataset included 46 NGS datasets from 41 patients, utilizing both Sanger and next-generation sequencing to the HIV-1 *pol* gene (SS+NGS).

The average pairwise Hamming distance of sequences in the quasispecies (referred to as “quasispecies diversity” later on) generated by SANTA-SIM (see Figure 2A-B) aligns well with empirical observations (Figure 2C-D). Our results exhibit temporal dynamics, showing low-diversity patterns in simulations with small generation numbers reminiscent of recent infections, and high quasispecies diversity in longer simulations resembling chronic HIV-1 infections. These are consistent with the observations of Shankarappa et al. [64], measuring sequence diversity within the *env* gene (∼600bp) using longitudinal samples from HIV-1 infected patients (see Figure 2C). The per-base Hamming distance (at the plateau) in chronic-like *in silico* sequence sets (IQR: 0.029-0.039) closely mirrors the empirical results of Shankarappa et al. [64] (IQR: 0.017-0.041) and is also very similar to the overall diversity of near-full-length sequences in the SGS-FULL dataset (IQR: 0.020-0.026) (Figure 2D). In our simulations, the genetic distance of the quasispecies consensus sequence from the HXB2 reference genome (referred to as “divergence from reference” later on) varied strongly with the subtype of the sequence used for the simulations and increases steadily but weakly with the duration of the simulations (Figure 2B).

**Figure 2:**
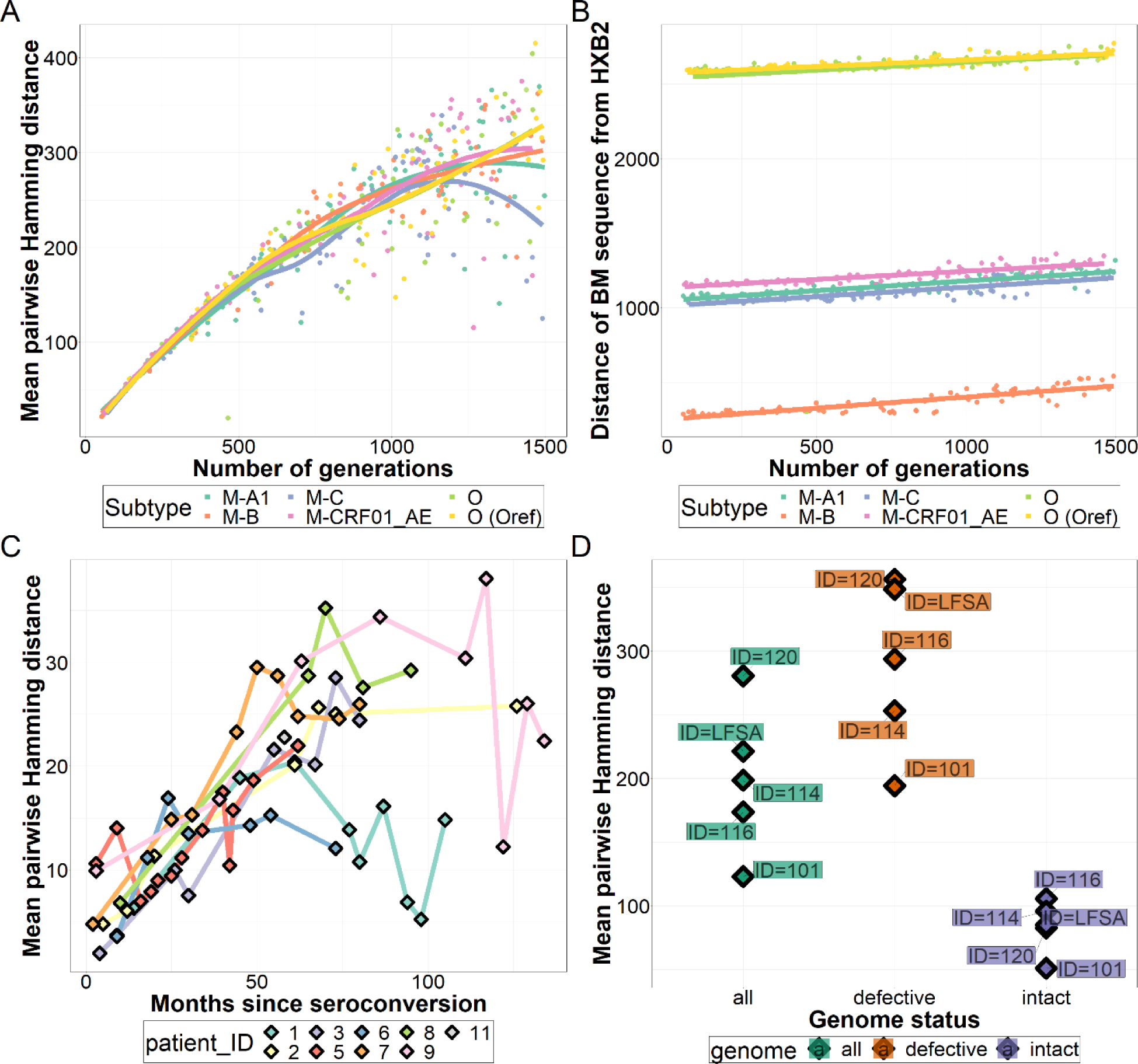
Within-host viral diversity of empirical and simulated quasispecies. Changes in A) quasispecies diversity and B) divergence from reference for all subtypes by the number of generations in the simulated datasets. C) Mean sequence diversity of the *env* gene throughout infection (for samples with 10 or more sequences) reported by Shankarappa et al [64]. D) Mean average Hamming distances for defective and intact sequence subpopulations and overall populations in the SGS-FULL dataset. Note that Hamming distance was not normalized for fragment length, which differed between the scenarios. Abbreviations: BM – benchmarking

### Quality of consensus genome assemblies and minority variant calls

Three out of the four pipelines exhibited failures (aborted runs). Specifically, viral-ngs showed eight failure events, V-Pipe only two, while shiver encountered errors in 25 out of 480 simulations. All assembly failures of shiver occurred during the random seed generation step of the *de novo* assembly process. This step requires sufficient overlapping reads to initiate contig generation, which explains why all errors manifested in low-coverage scenarios. Quasispecies diversity and distance from reference showed weak or no effect on failure events (logistic regression, odds=1.01, p=0.03 and odds=1.00, p=0.48, respectively).

We investigated the impact of variations in divergence from the HXB2 reference genome on the reliability of consensus genome assembly. In Table 2, we show the comparative performance of the four pipelines with two distinct sets of subtype/reference scenarios: in one set, the fixed-reference pipelines were used with the default HXB2 reference sequence to analyze the non-B datasets; in the other set, the reference sequence used by the fixed-reference pipelines (SmaltAlign, viral-ngs, V-Pipe) was selected to match the subtype/group of the sample (the subtype B scenario from the main analysis, which matches the default HXB2 reference sequence; and the additional group O scenario, where a group O reference sequence was used).

**Table 2:**
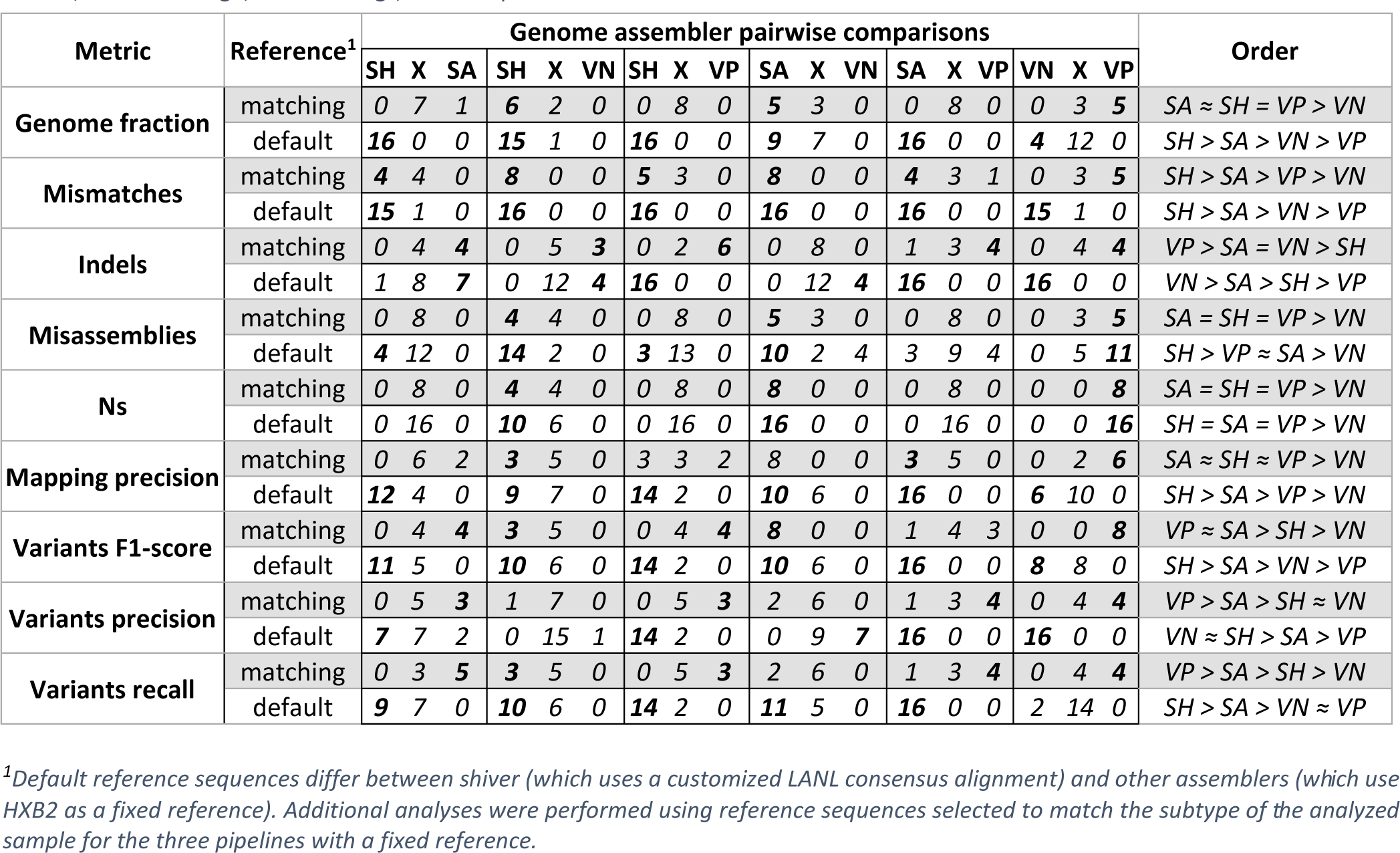
Comparison of genome assemblers based on quality metrics in the SIM dataset analyses. Quality metrics for genome assemblers were sed on the analyses that used the SIM dataset, and significant differences (fdr-adjusted p-value < 0.05) were determined through Wilcoxon d-rank tests on paired samples. The performance of assemblers was ordered based on the count of significant differences observed across simulated scenarios with relations categorized as equal (0, =), comparable (1-2, ≈) or differential (3+, >). Abbreviations: X – not significant, hiver, SA – SmaltAlign, VN – viral-ngs, VP – V-Pipe

Genome assemblers exhibited considerable differences in genome quality (Figures 3 and 4) and subsequent read mapping and variant calling (Figures 5) when the default reference sequence settings were applied. Apart from indel rates, shiver consistently outperformed other assemblers, achieving complete recovery, displaying the lowest mismatch rates, an absence of misassemblies and variant calling with high precision and recall irrespective of subtype, coverage, or contamination. The main reason behind this robust performance regardless of viral subtype is shiver’s unique feature to select the reference genome that is most similar to the *de novo* contigs for the imputation of missing regions.

**Figure 3:**
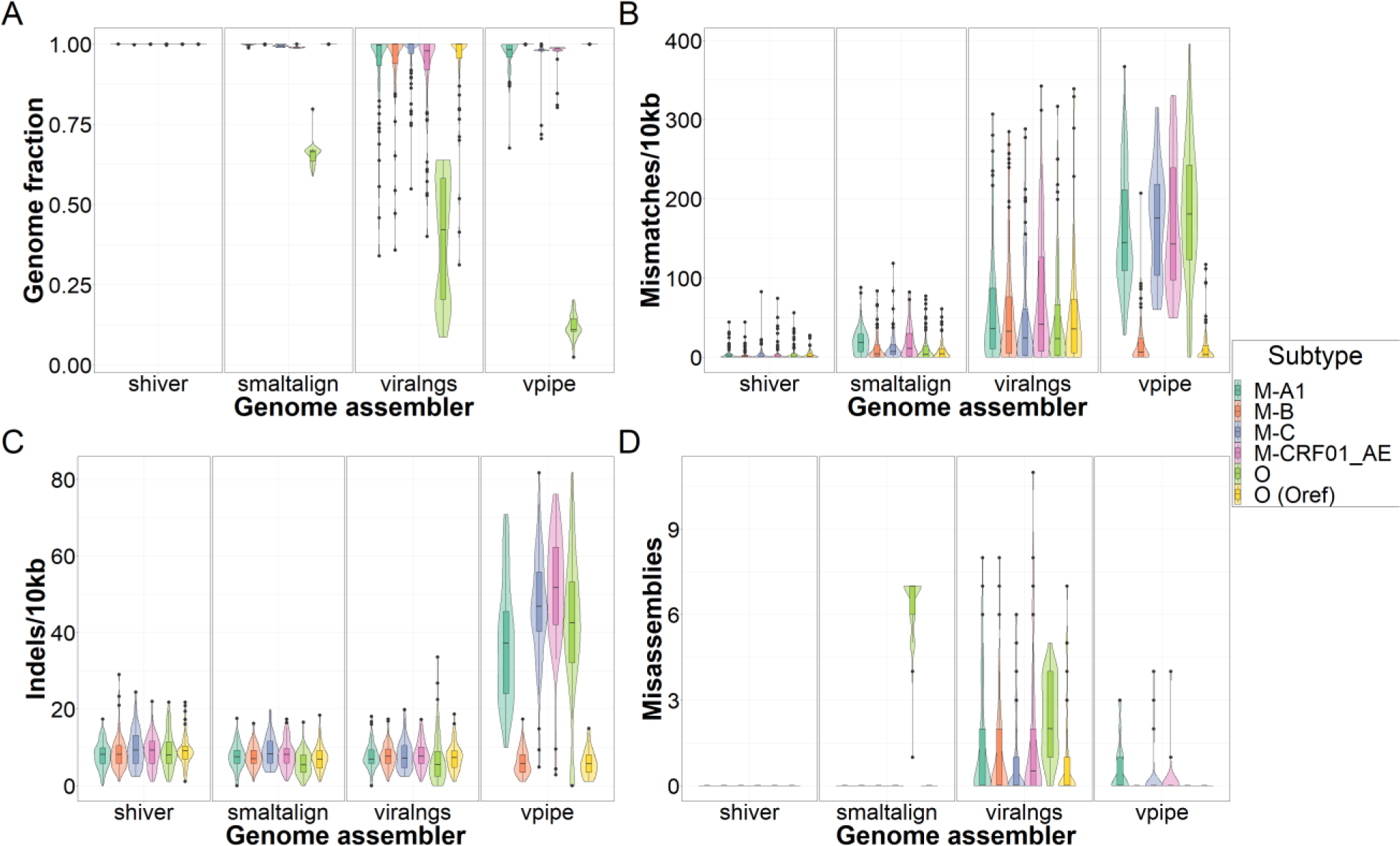
Assembly quality by viral subtype in the SIM dataset. Assemblers are compared based on A) the proportion of recovered positions in the genome, B) the rate of single nucleotide mismatches, C) the rate of small insertions and deletions (indels), and D) the number of misassemblies compared to the benchmarking sequence. The data shown include both coverage and contamination scenarios.

**Figure 4:**
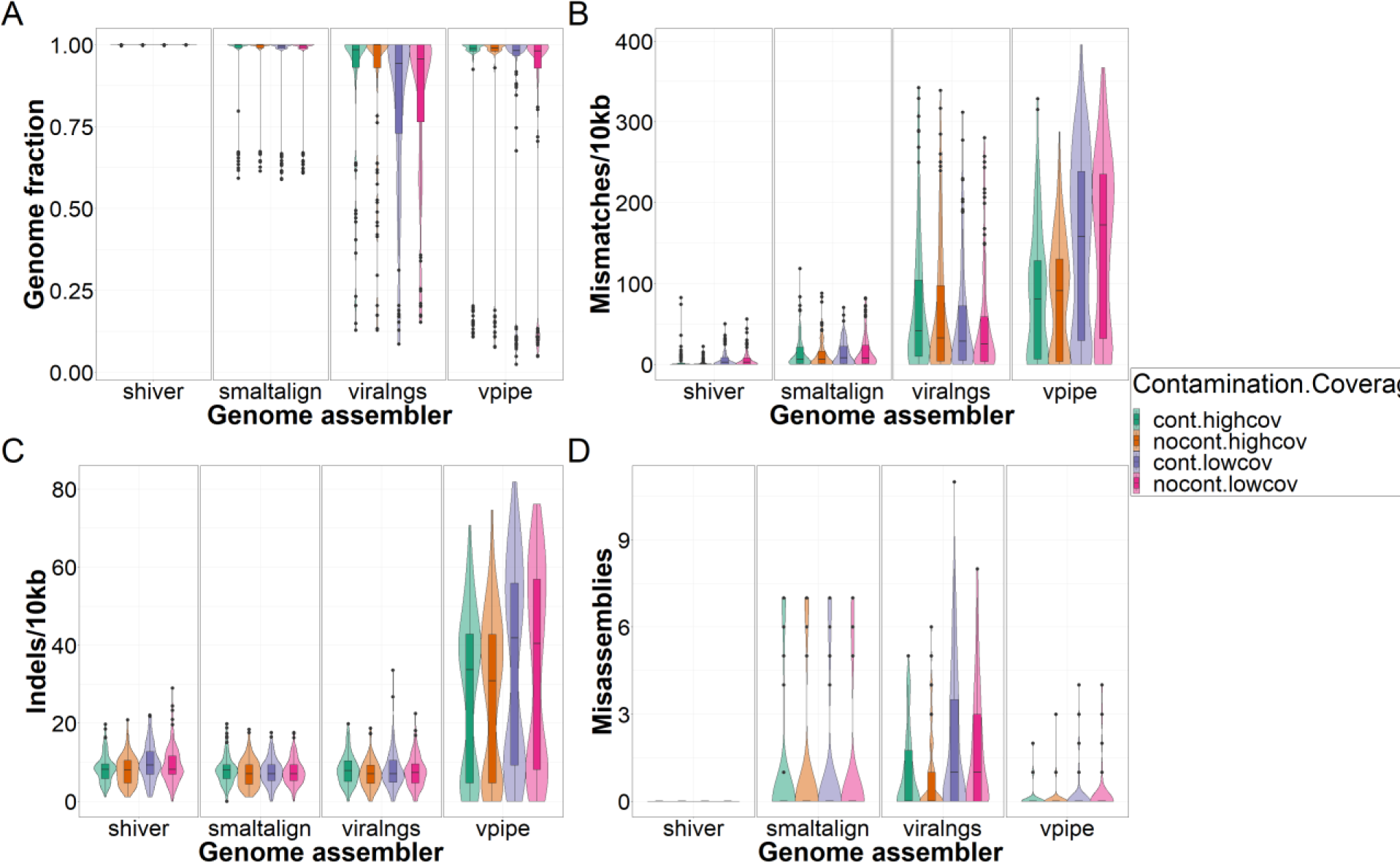
Assembly quality by scenario in the SIM dataset. Assemblers are compared based on A) genome fraction, B) mismatch rates, C) indel rates and D) the number of misassemblies compared to the benchmarking sequence. The data shown include simulation results with all virus types. Abbreviations: cont – contaminated, nocont – not contaminated, highcov – high coverage, lowcov – low coverage.

**Figure 5:**
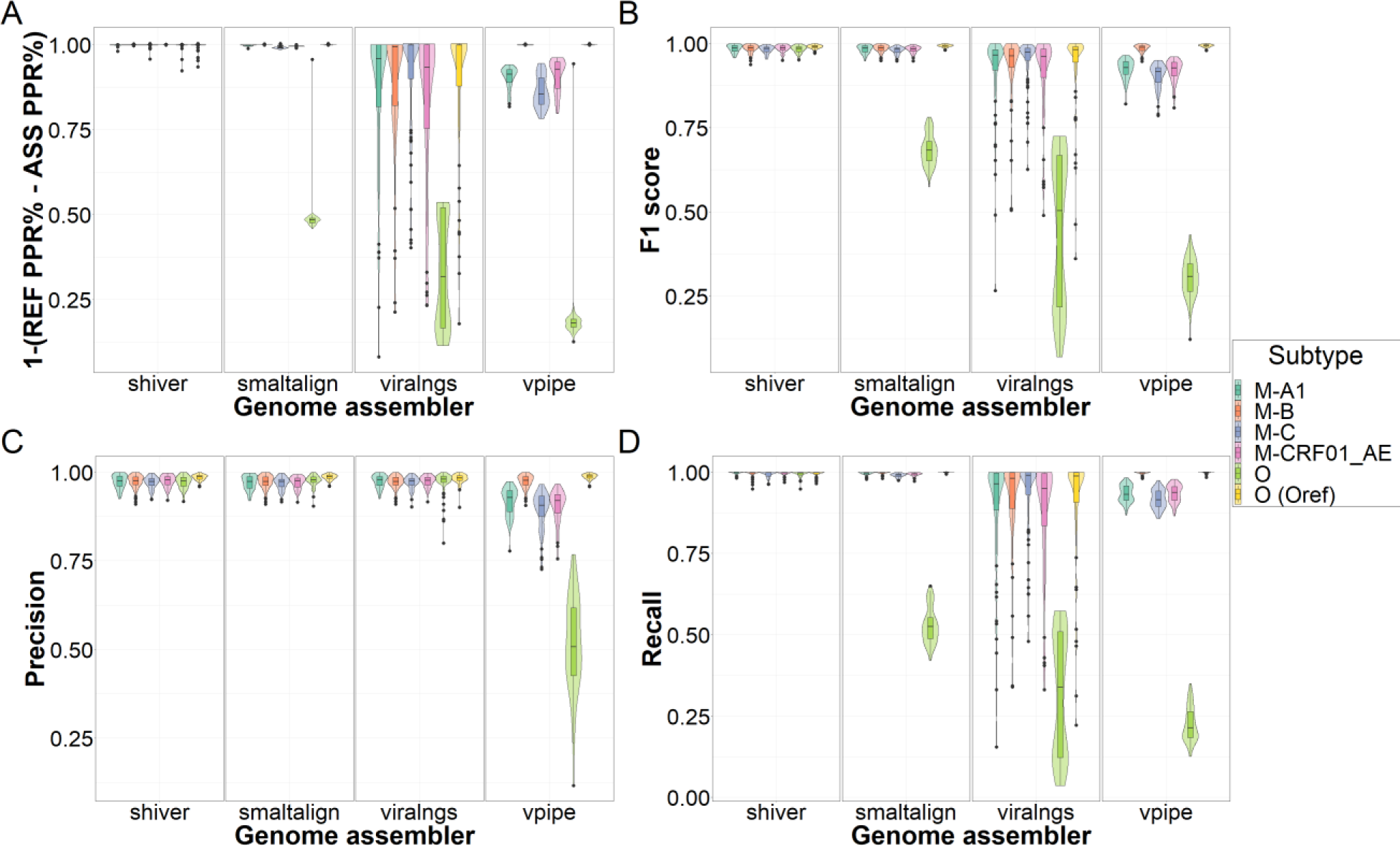
Read mapping and variant calling by viral subtype in the SIM dataset. Assemblers are compared based on A) the difference in the rate of properly paired reads between the assembly and the reference mapping, and B) the F1 score, C) the precision and D) the recall of minority variant calling compared to the benchmarking variant set. The data shown include both coverage and contamination scenarios. Abbreviations: REF - reference, ASS - assembly, PPR% - percentage of properly paired reads.

While their pairwise differences were statistically significant (Table 2), Smaltalign’s overall performance in most scenarios was comparable to that of shiver, with small numerical differences in all metrics. However, it exhibited a drop in quality metrics for group O samples, especially concerning genome fraction and misassemblies. This decrease in quality can be attributed to the assembler’s challenges in reconstructing certain genome segments in samples with high divergence from the fixed subtype B reference. In regions with low-quality reconstruction reads did not align to the assembly causing minority variants not being identified (drop in recall and F1 scores).

Genome assemblies generated by viral-ngs exhibited notably low indel rates and moderate mismatch rates. However, the reconstruction quality in some regions was suboptimal, leading to gaps with uncalled bases in the final assembly and a subsequent drop in read mapping precision and variant calling recall and F1 scores.

In samples with high divergence from reference, reference-based assembly can result in a biased loss of information due to the inability to map non-matching reads to the reference [40]. As anticipated, the reference-based assembler of V-Pipe was also significantly affected by divergence from the reference. This led to elevated mismatch and indel rates, a reduction in genome fraction, an increase in the number of misassemblies and lower variant calling recall and F1-scores in samples with highly divergent genomes. Unlike the other pipelines, V-Pipe predicted a considerable amount of false positive minority variants causing a drop in precision.

In cases where the reference sequence supplied for the fixed-reference pipelines matched the sample, we observed only minor but significant differences between assemblers (comparing them also to shiver). Notably, viral-ngs performed significantly worse compared to all other pipelines regarding genome fraction, misassemblies, uncalled bases and read mapping precision. Additionally, genome sequences produced by V-Pipe showed superior indel rate and variant calling metrics compared to all other genome assemblers when matching reference sequences were used. However, these differences carried through only weakly to later analysis steps. These results indicate that if the subtype (V-Pipe) or group (shiver, SmaltAlign, viral-ngs) of the analyzed sample is known and a reference sequence is selected accordingly for the genome assembly, each pipeline is capable of producing almost perfect consensus genomes from the simulated data.

We performed an additional analysis to investigate the pipeline-specific effects of sample properties and genome assembly quality on downstream analysis steps (Figure 6). First, we identified the order of analysis steps (quasispecies simulation, genome assembly, read mapping and variant calling) then matched each step with corresponding output metrics. After that, we conducted Spearman correlation tests on all variable pairs for each genome assembler (adjusting p-values using the Benjamini-Hochberg method) to reveal statistical associations between benchmarking metrics. Combining the correlation tests with the order of analysis steps enabled us to identify potential causal relationships between variables in adjacent analysis steps (because the order of steps is fixed, correlation between metrics obtained from subsequent analysis steps is likely to indicate the effect of the earlier step on the next step). The results of this analysis suggest that divergence from the reference sequence used for assembly had a considerable effect on genome completeness (Figure S1A), which had a negative impact on the precision of read mapping and the recall (and F1 scores) of minority variant detection (Figure S1B). Additionally, quasispecies diversity affected the mismatch and indel rates of genome assemblies (Figure S1C-D), and for V-Pipe and viral-ngs, and these metrics substantially altered subsequent steps of the analysis as well (Figure S1E). In the case of viral-ngs, a greater number of uncalled bases caused a drop in read mapping precision and minority variant calling (Figure S1F), while for V-Pipe, upstream analysis steps affected not only the recall but also the precision of minority variant calling.

**Figure 6:**
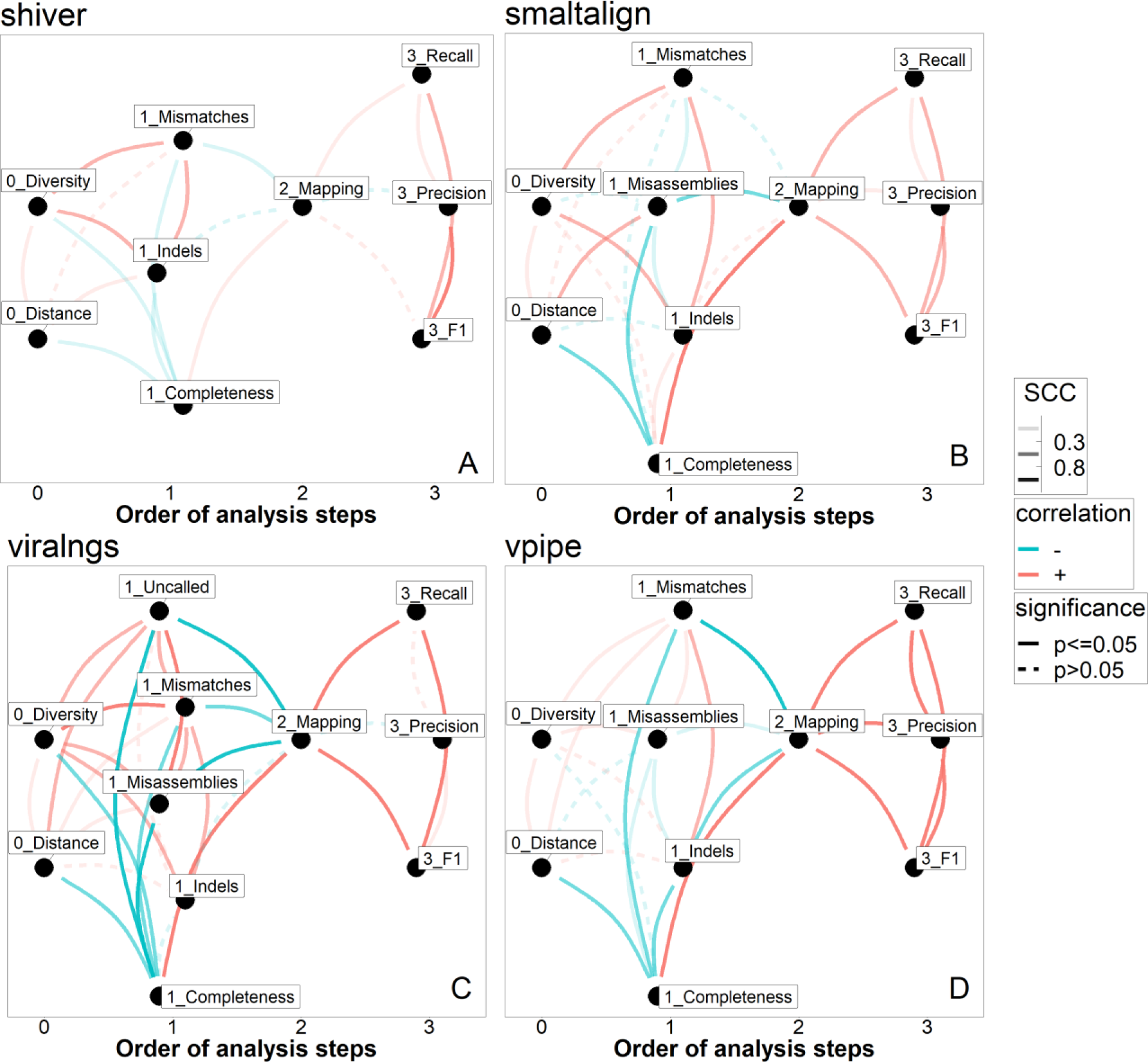
Correlative and potential causal relationships among benchmarking metrics. Undirected acyclic graphs for A) shiver, B) SmaltAlign, C) viral-ngs, and D) V-Pipe illustrating correlative (same analysis step) and potentially causal relationships (adjacent analysis steps) established through Pearson correlation tests between variables. Because the order of steps is fixed, correlation between metrics obtained from subsequent analysis steps is likely to indicate the effect of the earlier step on the next step. Order of analysis steps: 0 – quasispecies simulation, 1 – genome assembly, 2 – read mapping, 3 – variant calling. The scale of opacity of the edges reflects the absolute value of the correlation coefficient. Abbreviations: SCC – Spearman correlation coefficient.

For both the SGS-FULL and SS+NGS datasets, assembly quality and variant calling results were nearly equivalent among the assemblers (Supplementary Figures S2–5). The consistently reliable results achieved by all assemblers in this analysis are not surprising, given that the SGS-FULL dataset comprises samples from patients infected with subtype B viral populations only, and the SS+NGS scenario only assesses the quality of the short and high coverage pol regions of the near-full-length genome assemblies.

### Computational resource use

Maximum memory usage was similar across the examined pipelines, except for viral-ngs, which demonstrated lower memory requirements when dealing with low-coverage datasets (Figure 7A-B). However, we observed substantial variations in CPU time. In the SIM dataset analyses viral-ngs demonstrated the shortest runtime, closely followed by SmaltAlign and V-Pipe, while shiver required nearly an order of magnitude more runtime (Figure 7D). The runtime of all genome assemblers was influenced by genome coverage, and shiver, as the only pipeline that performs contaminant filtering, was particularly affected by contaminant reads. Furthermore, shiver lacks multithreading support (Figure 7C), resulting in even greater differences in elapsed real time compared to viral-ngs and SmaltAlign (Supplemetary Figure S6). Finally, in the analyses of empirical read sets, while shiver’s runtime was still longer compared to viral-ngs and Smalt-Align, V-Pipe required the longest CPU time to complete genome assembly (Figure 7E-F).

**Figure 7:**
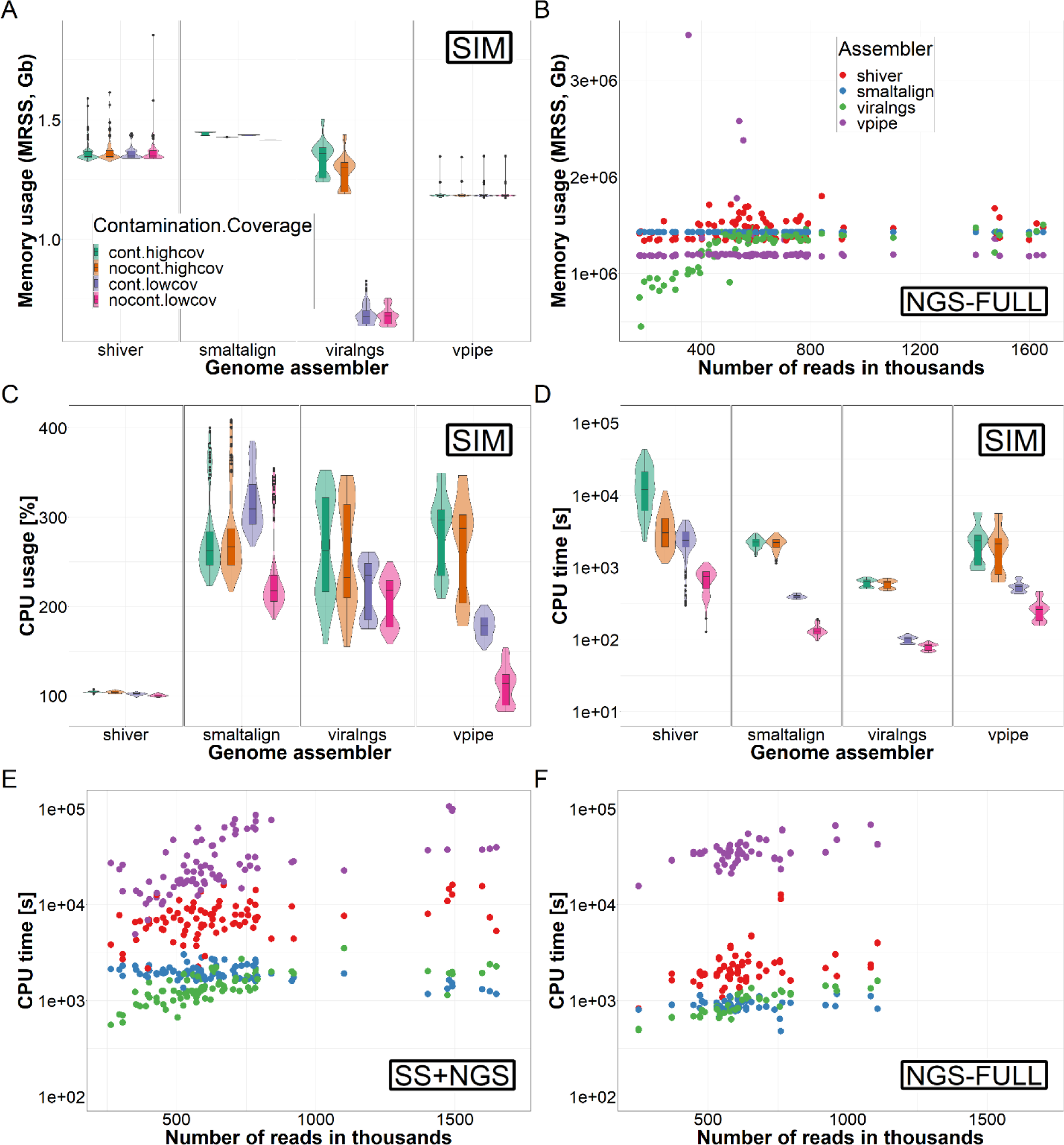
Runtime and memory usage by scenario. Comparison of genome assemblers based on computational needs. Maximum memory usage is compared in the A) SIM and B) NGS-FULL datasets, C) CPU usage in the SIM dataset, and CPU time in the D) SIM, E) NGS-FULL and F) SS+NGS datasets. Panels A, C and D stratify results according to coverage and contamination scenarios, panels B, E and F shows trends with varying dataset size (number of reads). Abbreviations: MRSS – maximum resident set size, cont – contaminated, nocont – not contaminated, highcov – high coverage, lowcov – low coverage.

Results of the benchmarking analyses for all three datasets are summarized in Figure 8 and Supplementary Figures S7–9.

**Figure 8.**
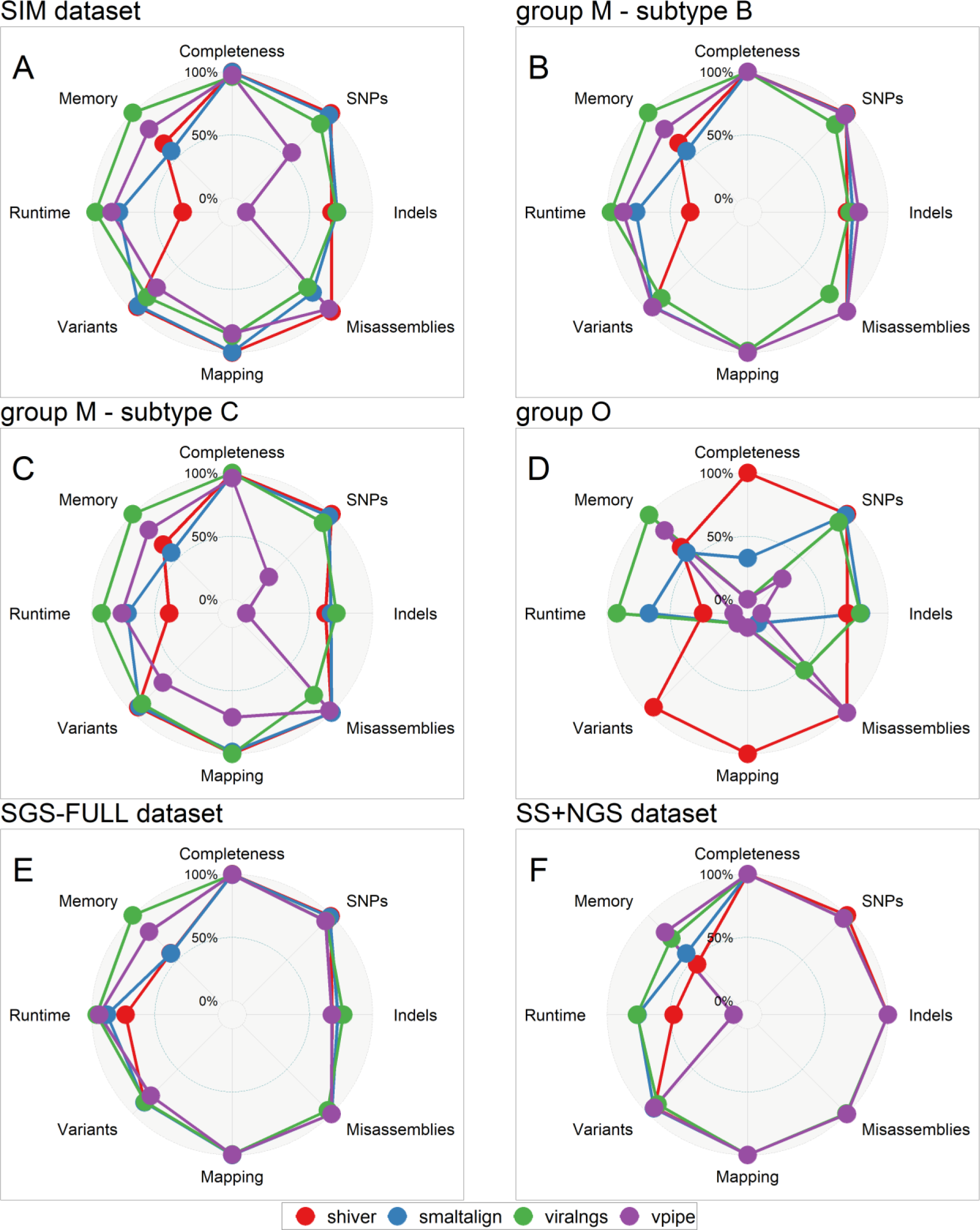
Multidimensional performance of genome assemblers. Benchmarking metrics are compared for A) the SIM dataset, in particular B) for subtype B, C) subtype C and D) group O results, and for E) the SGS-FULL and F) the SS+NGS datasets. For each metric the relative score between 100% and 0% is calculated using the following threshold values: Completeness – 100% and 50% median genome fraction, SNPs – 0 and 250 median mismatch rate/10kb, Indels – 0 and 25 median indel rate/10kb, Misassemblies – 0 and 5 mean misassemblies, Mapping – 100% and 50% median mapping precision, Variants – 1 and 0.75 median F1 scores, Runtime – 0h and 1h median user time and Memory – 1GB and 2GB median maximum resident set size.

### Ease of use

To utilize any of the examined genome assemblers, a basic understanding of the Linux command line and some knowledge of at least one environment management platform (Conda or Docker) are prerequisites. Multiple installation methods are available for all the pipelines, each accompanied by a comprehensive tutorial outlining the installation process and any dependency requirements. In our experience, due to the complexity of the tools, installation tends to require some troubleshooting unless the complete pipeline is available as a fixed containerized version. Among the four pipelines, SmaltAlign, viral-ngs and V-Pipe are integrated software that do not necessitate the installation of additional bioinformatic tools for basic functionality (V-Pipe offers the option of installing VICUNA for de novo reference construction). In contrast, the original version of shiver requires 7 additional tools for genome assembly. However, the containerized dshiver includes all dependencies (other than Docker itself).

Each pipeline is accompanied by a detailed user guide, including information on available parameters. All pipelines, except SmaltAlign, offer an extensive array of customizable parameters to fine-tune their performance for specific applications and some options to modify the workflow structure by selecting specific tools for various stages of the assembly algorithm. This makes SmaltAlign the easiest to use for typical use cases, but less customizable for new scenarios. All tools are capable of fully automated genome assembly, eliminating the need for additional programming skills or the manual processing of individual samples. Out of the four genome assemblers, shiver offers the most detailed output, including the assembled de novo contigs, position-specific depth and base frequencies, and separate consensus genomes with or without imputation from a reference sequence (and a drug resistance report in dshiver), which are not available in the other pipelines.

The simplified installation process, improved dependency handling, user-friendly automated workflow, and the detailed user guide included with dshiver enable the installation and use of this pipeline with basic computer user skills, on all main operating systems.

In terms of scalability, viral-ngs and V-Pipe stand out as the most convenient tools for large-scale analyses, offering mass importing, batch analysis, and advanced multithreading features as integral components of their Snakemake pipelines. In contrast, the other tools require some proficiency in shell scripting to perform analyses on multiple samples simultaneously.

## Discussion

Our analyses yielded two main insights. First, all four widely used viral NGS assembly pipelines can produce high-quality genome assemblies, which is a reassuring validation of the wealth of results that have been generated with these tools. Second, the only critical consideration is the appropriate choice of the reference sequence used for assembly. In the analysis of HIV samples, we observed relatively poor performance of some genome assemblers on samples with high distance from the fixed subtype B (HXB2) reference sequence (group O simulations). We note, however, that in the majority of applications group M samples will be analyzed, and the observed suboptimal performance with divergent samples can be easily improved by using a matched reference sequence, either by implementing automatic selection (as in the shiver pipeline), or by supplying a matching reference sequence for the current pipelines. While it is still common practice to employ the HXB2 sequence as a reference for HIV-1 genome assembly [65–69], selecting a more suitable reference sequence is a viable option even without the modification of the genome assembler pipeline. To guide the selection, the predominant viral subtype can be determined by HIV subtyping tools such as REGA [63] or COMET [70], using either Sanger sequencing data (if available from the same sample) or the result of the first round of genome assembly.

Our work is subject to limitations. While each investigated pipeline involves a large number of variable parameters that may influence performance metrics, we used fixed parameter sets, as constraints of time and computational resources restricted the exploration of further combinatorial dimensions in the parameter space. However, expert customization can only improve the performance of the pipelines and can therefore not affect the main finding of good performance by all four pipelines in the key quality metrics. Furthermore, the use of fixed (default) parameterization is a scenario that reflects most closely the use case of NGS analyses by non-specialists.

We employed HIV-1 data as a test case to assess genome assemblers. HIV-1 is one of the most extensively researched and medically significant viruses, demonstrating high levels of diversity both within and between hosts. However, none of the examined pipelines are specifically designed for, or restricted to HIV-1, and they can be readily adapted to other viruses. Our results emphasize the need to select a matching reference sequence for assembly, especially for other viruses, like hepatitis C virus where genetic distances tend to be larger than those observed for HIV [71]. Furthermore, the contaminant filtering feature of shiver may be more relevant if the samples contain phylogenetically closer contaminant reads (and may have to be used cautiously for viruses that tend to pick up unique human genomic fragments, like hepatitis E virus [72,73]).

Finally, we note that while sequencing read simulators aim to replicate various characteristics of sequencing data and biases caused by laboratory protocols [74], our chosen tool, ART, like other read simulators, has limitations in addressing some factors that may have impacted our analysis. These include biased amplification due to primer mismatches [75], which can cause certain haplotypes to appear more or less frequently in the read data, possibly reducing the performance of all assemblers; and variation in sequencing depth across different genomic regions, resulting in low coverage regions, a phenomenon often observed in empirical NGS results of HIV-infected samples [76,77]. The presence of these and other unidentified factors may lead to disparities between *in silico* and empirical read sets. Such disparities may have contributed to the observed differences in runtime performance between the SIM and the NGS-FULL and SS+NGS datasets.

In summary, our analysis addressed a gap in current research by benchmarking state-of-the-art genome assembly pipelines for small viruses. Reassuringly, all four widely used pipelines can perform well, although our results highlight some caveats, specific strengths of individual pipelines, and also differences in practical usability.

## Key points

All four pipelines (shiver, SmaltAlign, viral-ngs and V-Pipe) can perform well in terms of quality metrics; however, the reference sequence needs to be adjusted to closely match the sample data for viral-ngs and V-Pipe. Differences in user-friendliness and runtime may guide the choice of the pipeline in a particular setting. The new Dockerized version of shiver offers ease of use in addition to the accuracy and robustness of the original pipeline.

## Data availability

Sanger sequence data used in this study have been submitted to GenBank at https://www.ncbi.nlm.nih.gov/nucleotide/ under accession numbers MK213294, MK213306, MK236513,

MK236525, MK250657, MK250672, MK250680, PP313557-PP313598, PP333487-PP333522 and sequencing reads to the Sequence Read Archive at https://www.ncbi.nlm.nih.gov/sra under BioProject accession number PRJNA1078284 and BioSample accession numbers SAMN39993709-SAMN39993754 and have been tagged to be released within 2 years. The use of these data for the purposes of the study has been approved by the Scientific and Research Ethics Committee of the Medical Research Council, Budapest (reference number: BM/18124-1/2023).

## Funding

This work was supported by the National Research, Development and Innovation Office in Hungary (RRF- 2.3.1-21-2022-00006) as a part of the National Laboratory for Health Security.

## Acknowledgements

We are grateful to Prof. Niko Beerenwinkel and Lara Fuhrmann for useful discussions on the concept and early results of our analyses. We disclose the use of ChatGPT-3.5 solely to identify grammatical mistakes and improve text flow during manuscript preparation.

## Disclosure of potential conflicts of interest

K.J.M. has received travel grants and honoraria from Gilead Sciences, Roche Diagnostics, GlaxoSmithKline, Merck Sharp & Dohme, Bristol-Myers Squibb, ViiV and Abbott; and the University of Zurich received research grants from Gilead Science, Novartis, Roche, and Merck Sharp & Dohme for studies that K.J.M. serves as principal investigator for, and advisory board honoraria from Gilead Sciences and ViiV. R.D.K. received grants from Gilead Sciences, the National Institutes of Health, and the Swiss National Science Foundation (all to institution).

## Supplementary information: Next-generation sequencing protocol

A next-generation sequencing protocol was developed for the amplification of near full-length HIV-1 genome and short-read sequencing based on previous publications [1]. Viral RNA was extracted from 300 or 280 µl EDTA-treated plasma using NucliSENS miniMAG nucleic acid purification system and specific magnetic extraction reagents (bioMérieux) or QIAamp Viral RNA Mini kit (Qiagen) as per manufacturers’ instructions. Purified viral RNA was stored at -80°C or immediately reverse transcribed into cDNA applying a PrimeScript High Fidelity RT-PCR kit (Takara Bio) 2 step RT-PCR protocol with oligo dT or gene specific primers. Nearly full-length HIV-1 genomes were amplified in four overlapping fragments in nested PCR using MyFi Mix (2x) (Meridian Bioscience). Composition of reaction mixtures, thermal cycling conditions and PCR primers applied in reverse transcription and nested-PCR are listed in Table S1-S3. The quality of amplicons was verified by agarose gel electrophoresis. Four PCR amplicons of each sample were purified using AMPure XP beads (Beckman Coulter) and were pooled in equimolar amounts by using Qubit dsDNA HS Assay kit and Qubit 4 Fluorometer (Invitrogen). NGS library preparation was performed following the manufacturer’s protocol of Nextera XT Library Preparation and Nextera XT Index kit v2 (Illumina). All quantification steps were implemented using Qubit dsDNA HS Assay kit during library preparation. The size distribution of libraries was assessed using High Sensitivity D1000 Reagents and ScreenTape on TapeStation 2200 system (Agilent Technologies). Paired-end reads were sequenced with MiSeq Reagent v2 kit (300 cycles) on MiSeq instrument (Illumina). To evaluate the effects of nucleic acid extraction method, presence / absence of carrier RNA (QIAamp Viral RNA Mini kit), reverse transcription with oligo dT / gene specific primers, and one round / two round PCR on sequencing output, parallel testing of selected samples was performed, resulting in 46 NGS datasets from 41 patients.

### Supplementary tables: specifications of the reverse transcription and nested PCR protocol

**Table S2:** Composition of reverse transcription and nested PCR reaction mixtures

| Reaction | Reagent | Amount (μl) | Total amount (μl) |
| --- | --- | --- | --- |
| Template denaturation and primer annealing (M1) | dNTP Mixture (10 mM each) | 1 | 10 |
|  | oligo dT (2,5 μM) / gene specific primer (10 μM) | 1 |  |
|  | template RNA | 5 |  |
|  | RNase-free dH<sub>2</sub>O | 3 |  |
| Reverse transcription | denatured and annealed reaction mixture (M1) | 10 | 20 |
|  | 5x PrimeScript Buffer | 4 |  |
|  | RNase Inhibitor (40U/μl) | 0.5 |  |
|  | PrimeScript RTase | 0.5 |  |
|  | RNase-free dH<sub>2</sub>O | 5 |  |
| First round PCR | MyFi Mix, 2x | 12.5 | 25 |
|  | Forward primer (10 μM) | 1 |  |
|  | Reverse primer (10 μM) | 1 |  |
|  | Nuclease-free dH<sub>2</sub>O | 5.5 |  |
|  | cDNS | 5 |  |
| Second round PCR | MyFi Mix, 2x | 12.5 | 25 |
|  | Forward primer (10 μM) | 1 |  |
|  | Reverse primer (10 μM) | 1 |  |
|  | Nuclease-free dH<sub>2</sub>O | 8.5 |  |
|  | First round PCR product | 2 |  |

**Table S3:**
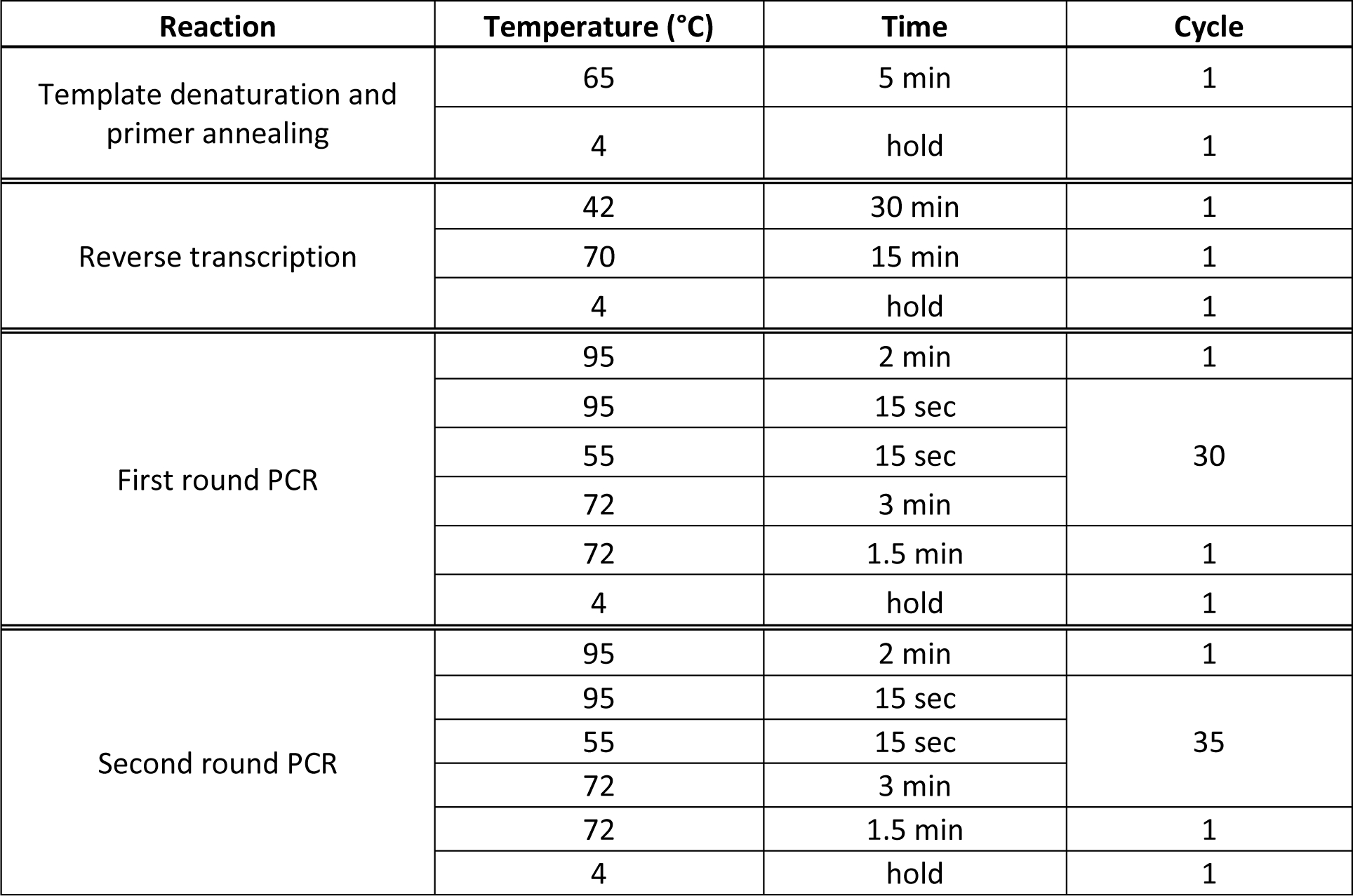
Cycling conditions of reverse transcription and nested PCR

**Table S4:** Primers used in reverse transcription and nested PCR

| Reaction | Amplicon 1 |  | Amplicon 2 |  | Amplicon 3 |  | Amplicon 4 |  |
| --- | --- | --- | --- | --- | --- | --- | --- | --- |
|  | Forward | Reverse | Forward | Reverse | Forward | Reverse | Forward | Reverse |
| Reverse transcription | - | GAG-RT-1R | - | oligo dT/<br>RT-IN-1R/<br>Pan-HIV-1_2R | - | oligo dT/<br>IN-ENV-1R | - | oligo dT/<br>ENV-NEF-1R |
| First round PCR | GAG-RT-1F | GAG-RT-1R | RT-IN-1F | RT-IN-1R/<br>Pan-HIV-1_2R | Pan-HIV-1_3F | IN-ENV-1R | ENV-NEF-1F | ENV-NEF-1R |
| Second round PCR | GAG-RT-2F | GAG-RT-2R | RT-IN-2F | RT-IN-2R/<br>Pan-HIV-1_2R | Pan-HIV-1_3F | IN-ENV-2R | ENV-NEF-2F | ENV-NEF-2R |
| Reference for primers [1,2] |  |  |  |  |  |  |  |  |

### Supplementary figures: Correlation of genome assembly metrics

**Figure S1:**
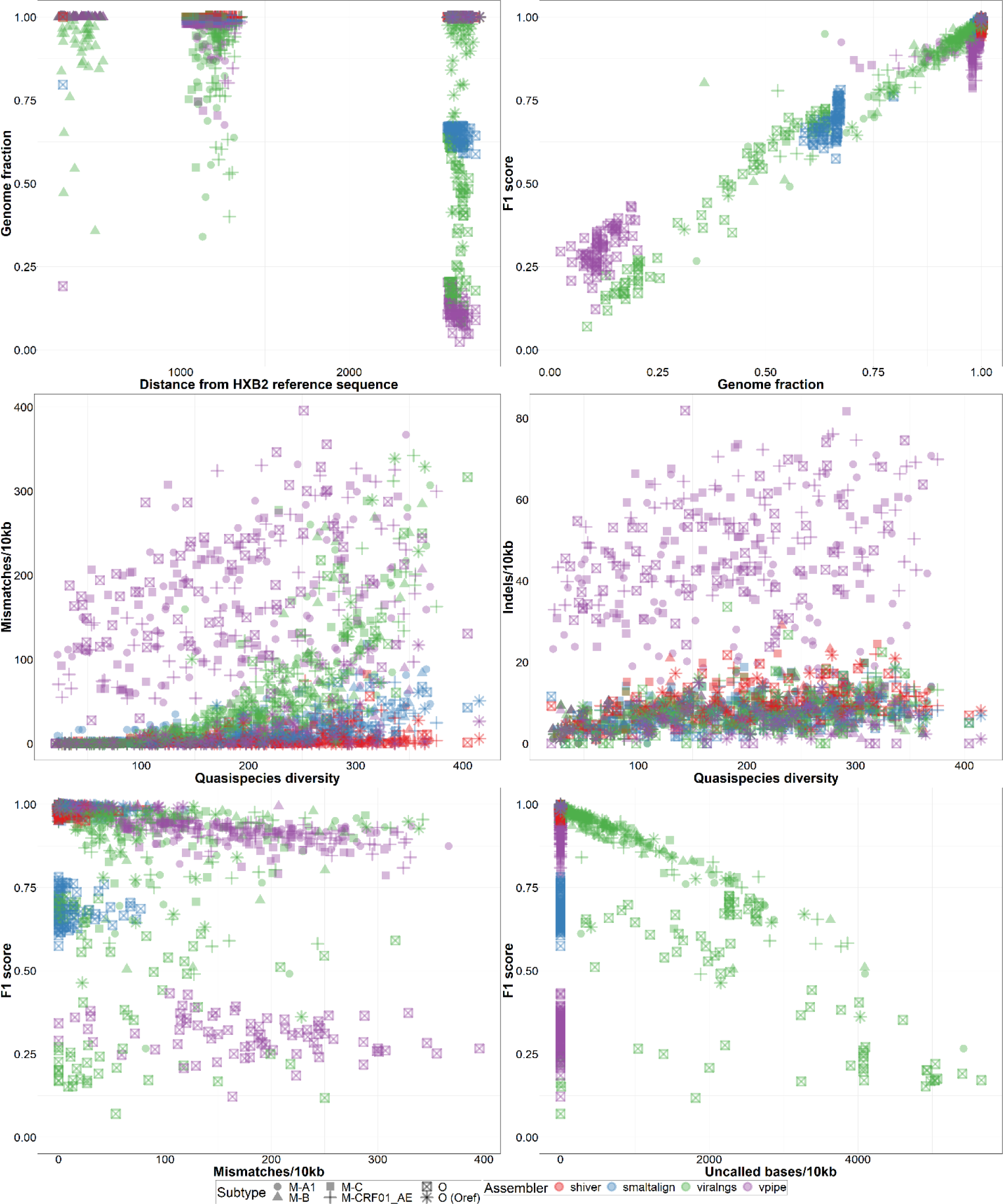
Correlation of genome assembly metrics in the SIM dataset. The panels show the relationship between A) distance from reference and genome fraction, B) genome fraction and variant calling F1 scores, C) quasispecies diversity and mismatch rates, D) quasispecies diversity and indel rates, E) mismatch rates and F1 scores and F) uncalled base rate and F1 scores. Points are grouped according to the genome assembler and the subtype of the sample.

### Supplementary figures: Quality of genome assemblies using the SGS-FULL dataset

**Figure S2:**
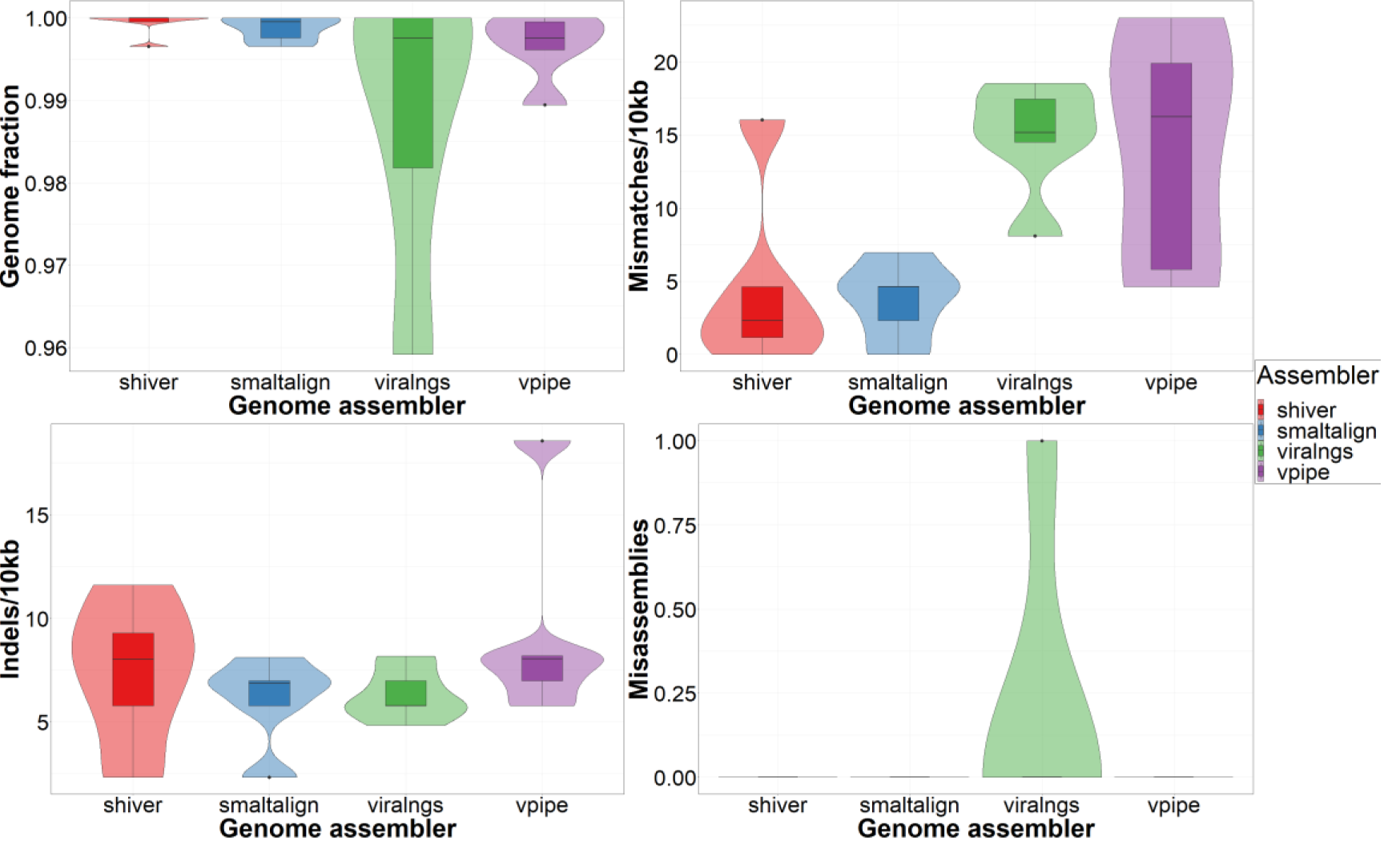
Assembly quality in the SGS dataset. Assemblers are compared based on A) the proportion of recovered positions in the genome, B) the rate of single nucleotide mismatches, C) the rate of small insertions and deletions (indels), and D) the number of misassemblies compared to the benchmarking sequence.

**Figure S3:**
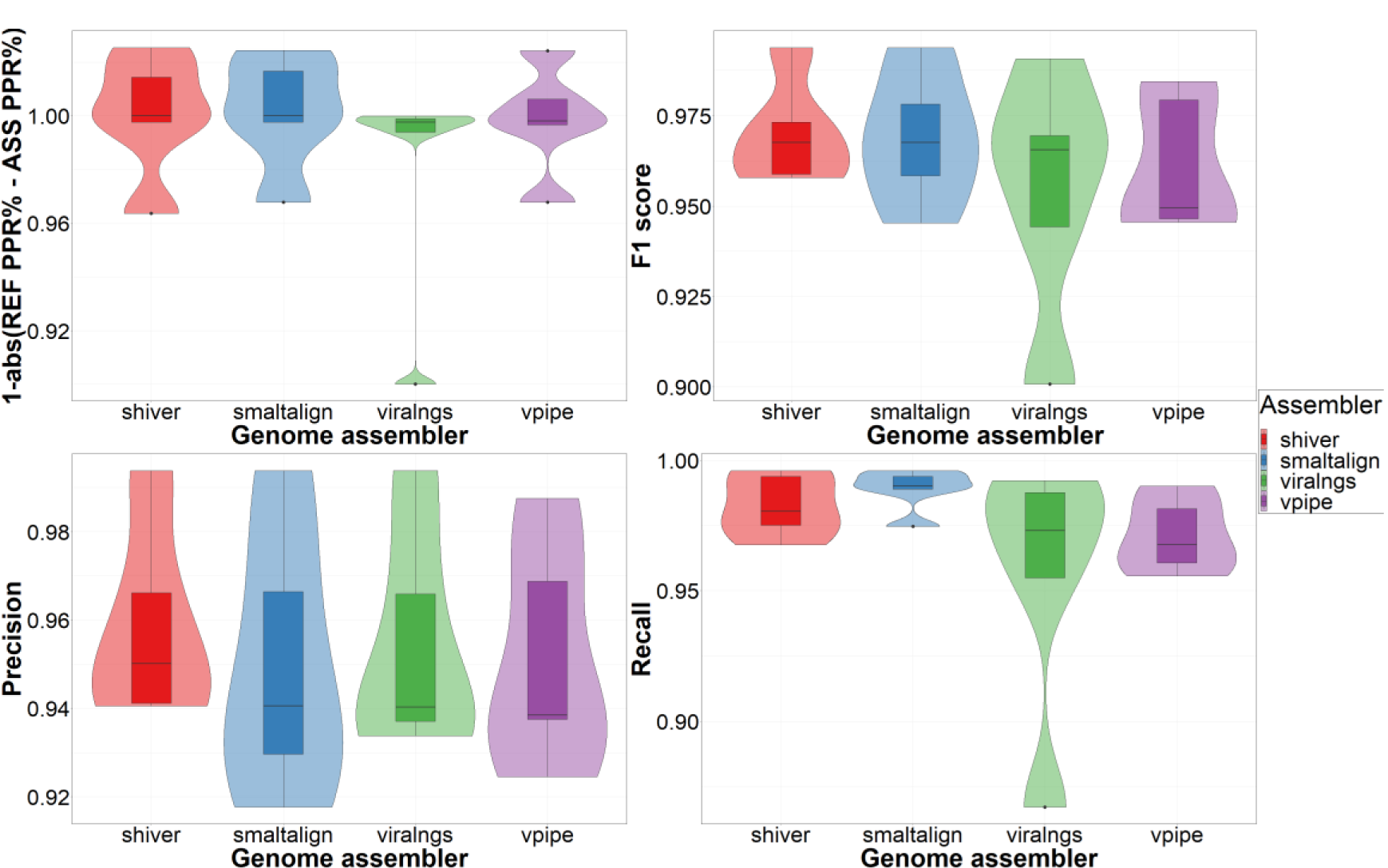
Read mapping and variant calling in the SGS-FULL dataset. Assemblers are compared based on A) the difference in the rate of properly paired reads between the assembly and the reference mapping, and B) the F1 score, C) the precision and D) the recall of minority variant calling compared to the benchmarking variant set. Abbreviations: REF - reference, ASS - assembly, PPR% - percentage of properly paired reads.

### Supplementary figures: Quality of genome assemblies using the SS+NGS dataset

**Figure S4:**
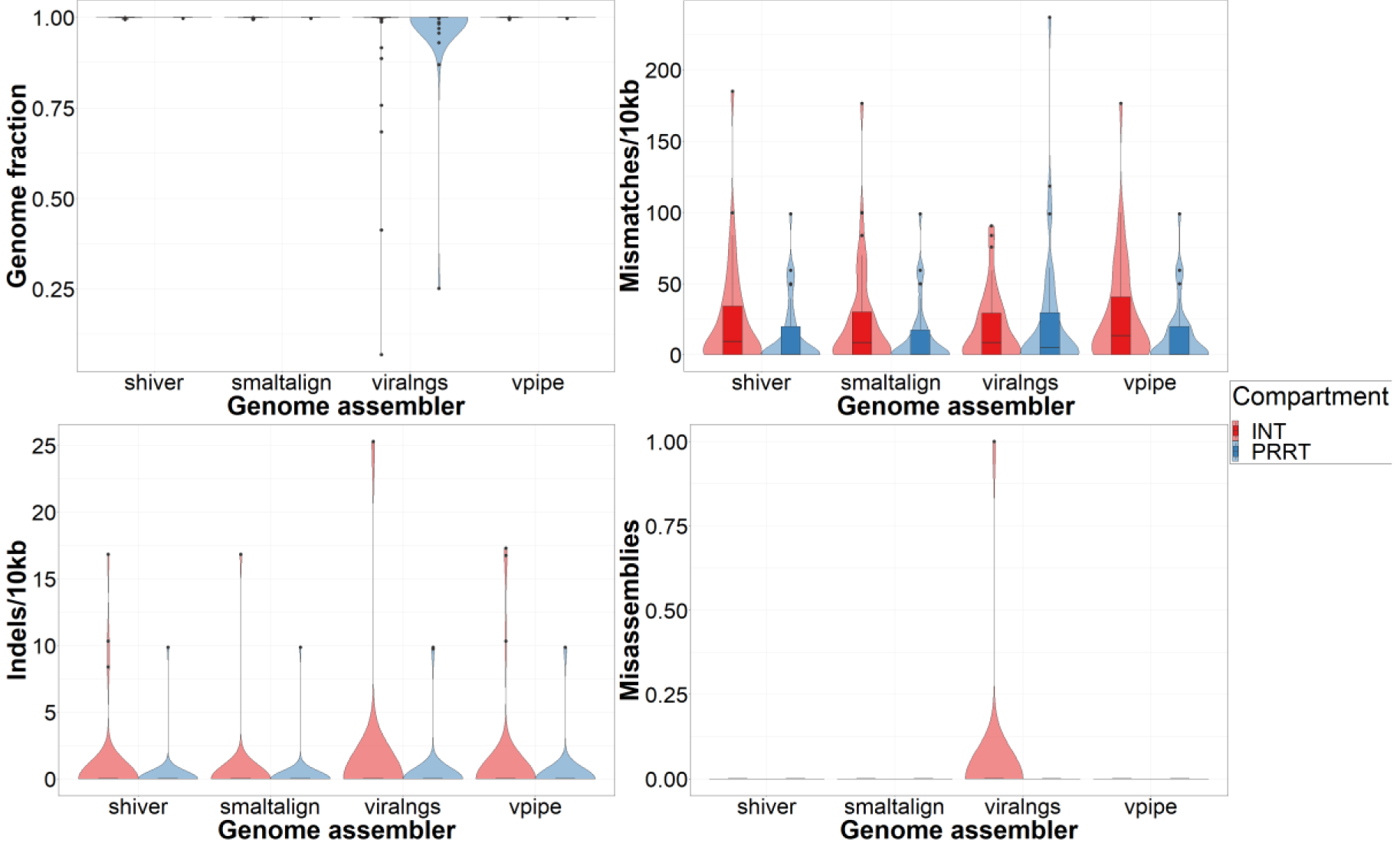
Assembly quality in the SS+NGS dataset. Assemblers are compared based on A) the proportion of recovered positions in the genome, B) the rate of single nucleotide mismatches, C) the rate of small insertions and deletions (indels), and D) the number of misassemblies compared to the benchmarking sequence.

**Figure S5:**
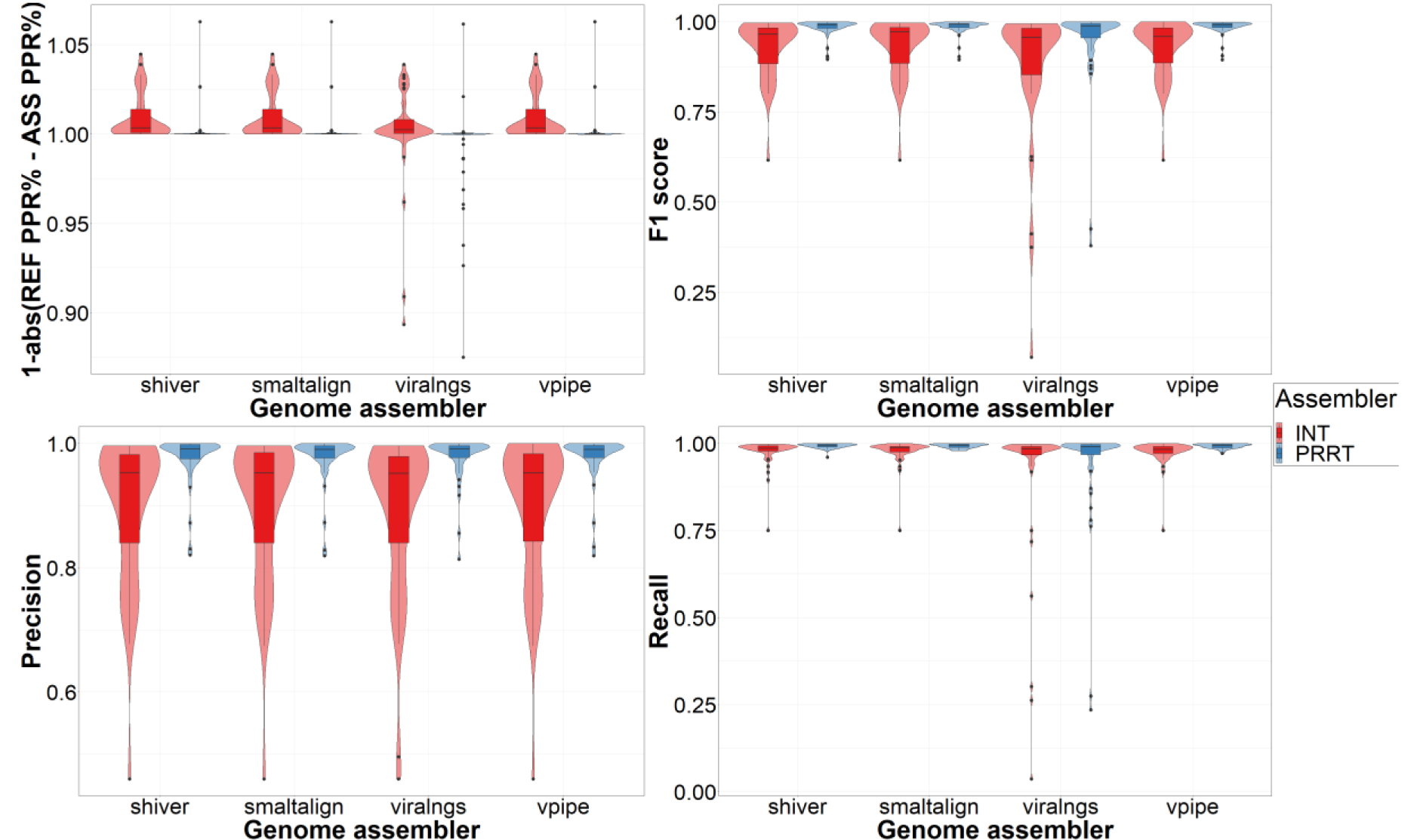
Read mapping and variant calling in the SS+NGS dataset. Assemblers are compared based on A) the difference in the rate of properly paired reads between the assembly and the reference mapping, and B) the F1 score, C) the precision and D) the recall of minority variant calling compared to the benchmarking variant set. Abbreviations: REF - reference, ASS - assembly, PPR% - percentage of properly paired reads.

### Supplementary figures: Comparison of real time requirements of the examined pipelines

**Figure S6:**
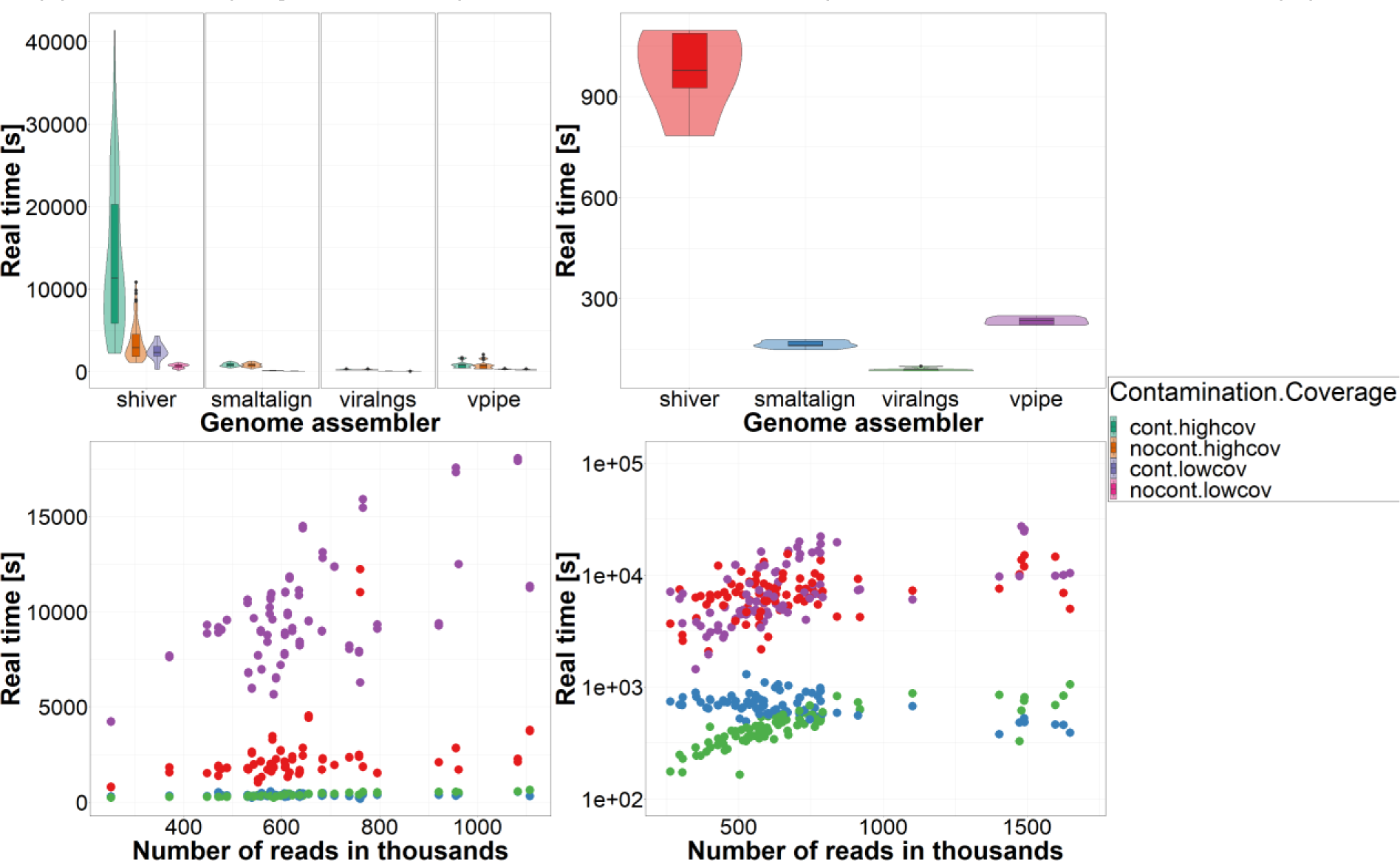
Comparison of elapsed time between assemblers. Comparison of genome assemblers based on runtime (real or elapsed time) in the A) SIM, B) SGS-FULL, C) SS+NGS and D) NGS-FULL datasets. Panel A stratifies results according to coverage and contamination scenarios and panels C and D show trends with varying dataset size (number of reads). Abbreviations: cont – contaminated, nocont – not contaminated, highcov – high coverage, lowcov – low coverage.

### Supplementary figures: Multivariate comparison of pipelines in two subtype scenarios

**Figure S7:**
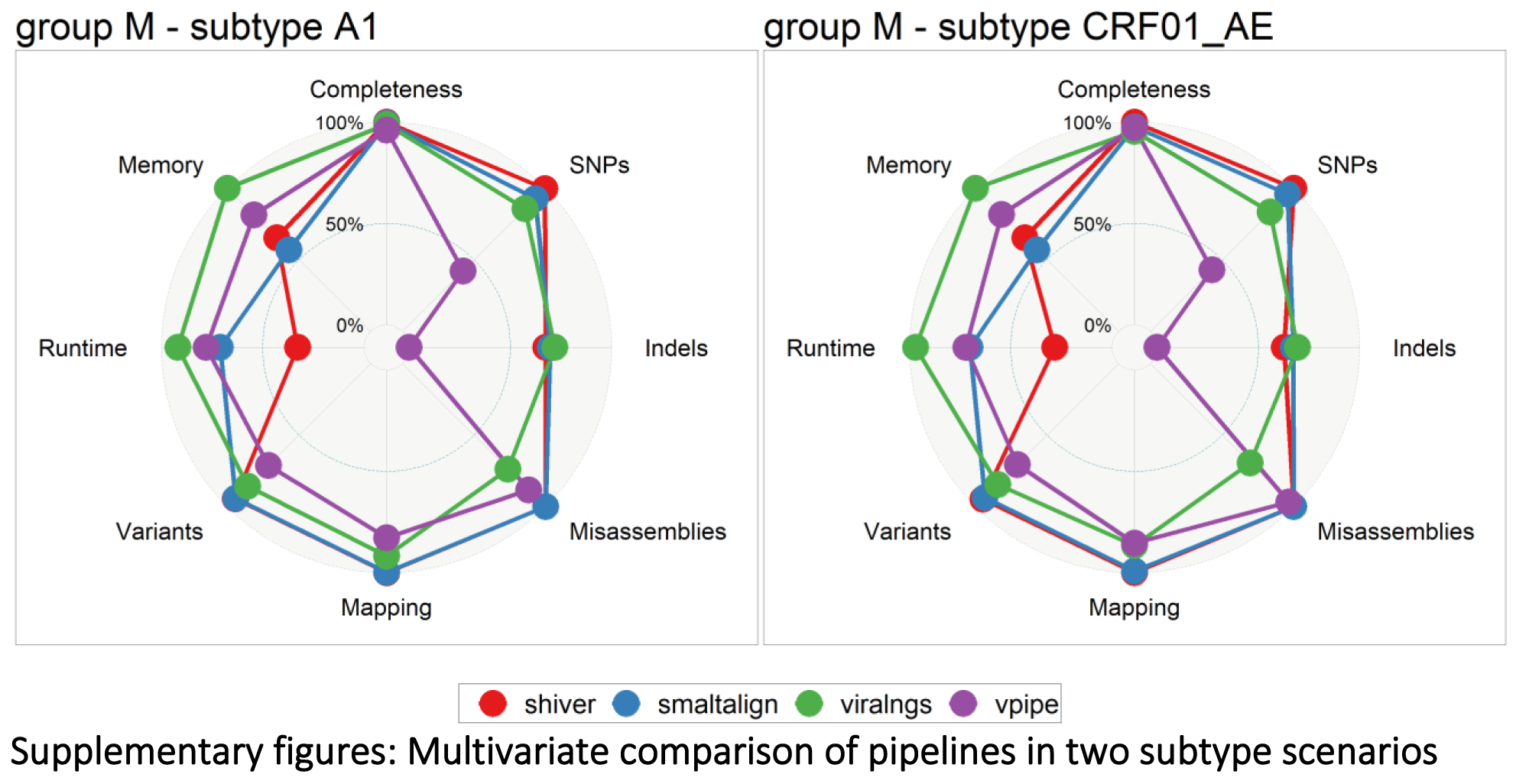
Multidimensional performance of genome assemblers in the subtype A1 and CRF01_AE scenarios. Benchmarking metrics are compared for A) the subtype A and B) for the CRF01_AE scenarios of the SIM dataset. For each metric the relative score between 100% and 0% is calculated using the following threshold values: Completeness – 100% and 50% median genome fraction, SNPs – 0 and 250 median mismatch rate/10kb, Indels – 0 and 25 median indel rate/10kb, Misassemblies – 0 and 5 mean misassemblies, Mapping – 100% and 50% median mapping precision, Variants – 1 and 0.75 median F1 scores, Runtime – 0h and 1h median user time and Memory – 1GB and 2GB median maximum resident set size.

### Supplementary figures: Multivariate comparison of pipelines in the combination of coverage and contamination scenarios

**Figure S8:**
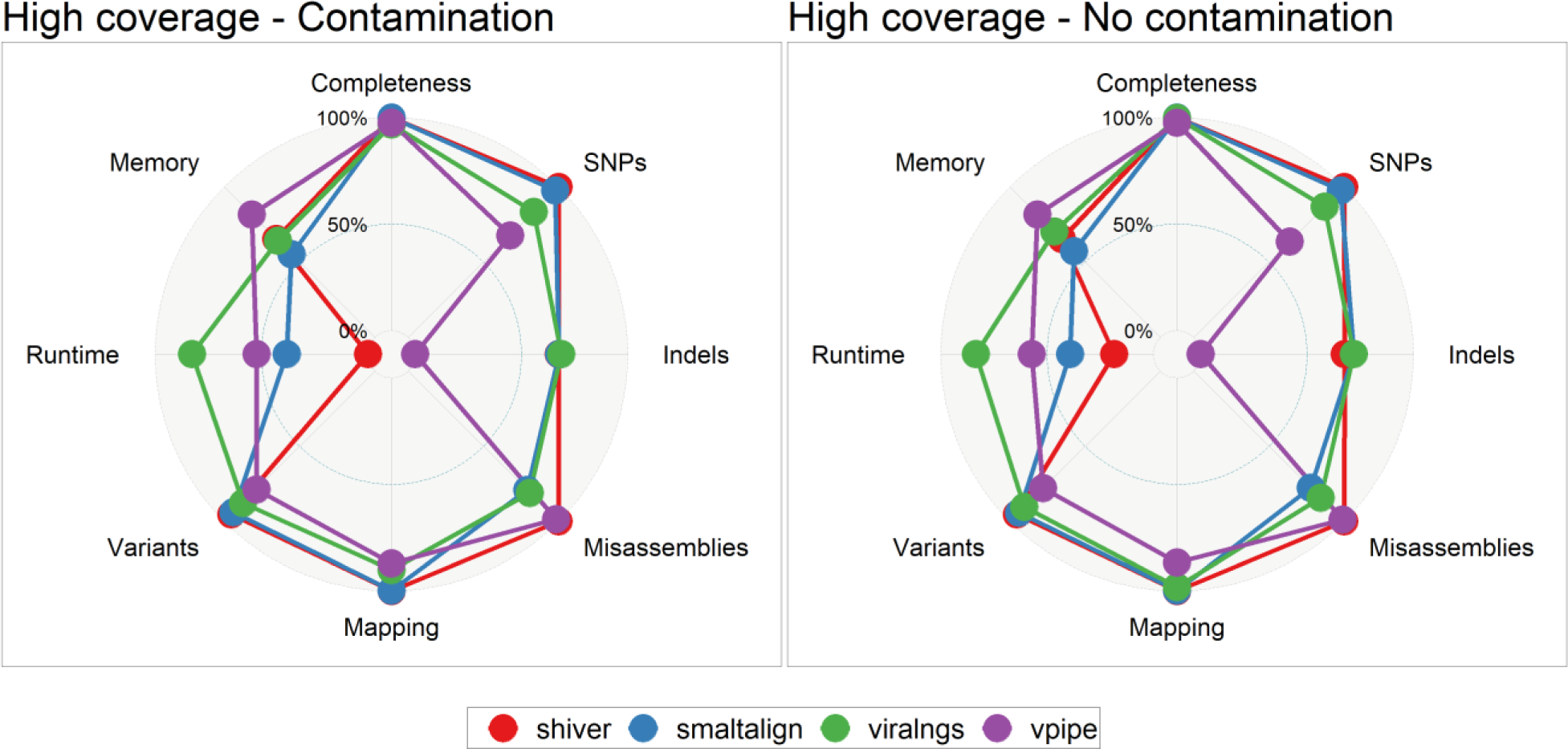
Multidimensional performance of genome assemblers in high coverage scenarios. Benchmarking metrics are compared for A) the high coverage – contamination and B) for the high coverage – no contamination scenarios of the SIM dataset. For an explanation on the scaling of the different axes see the figure caption of Figure S6.

**Figure S9:**
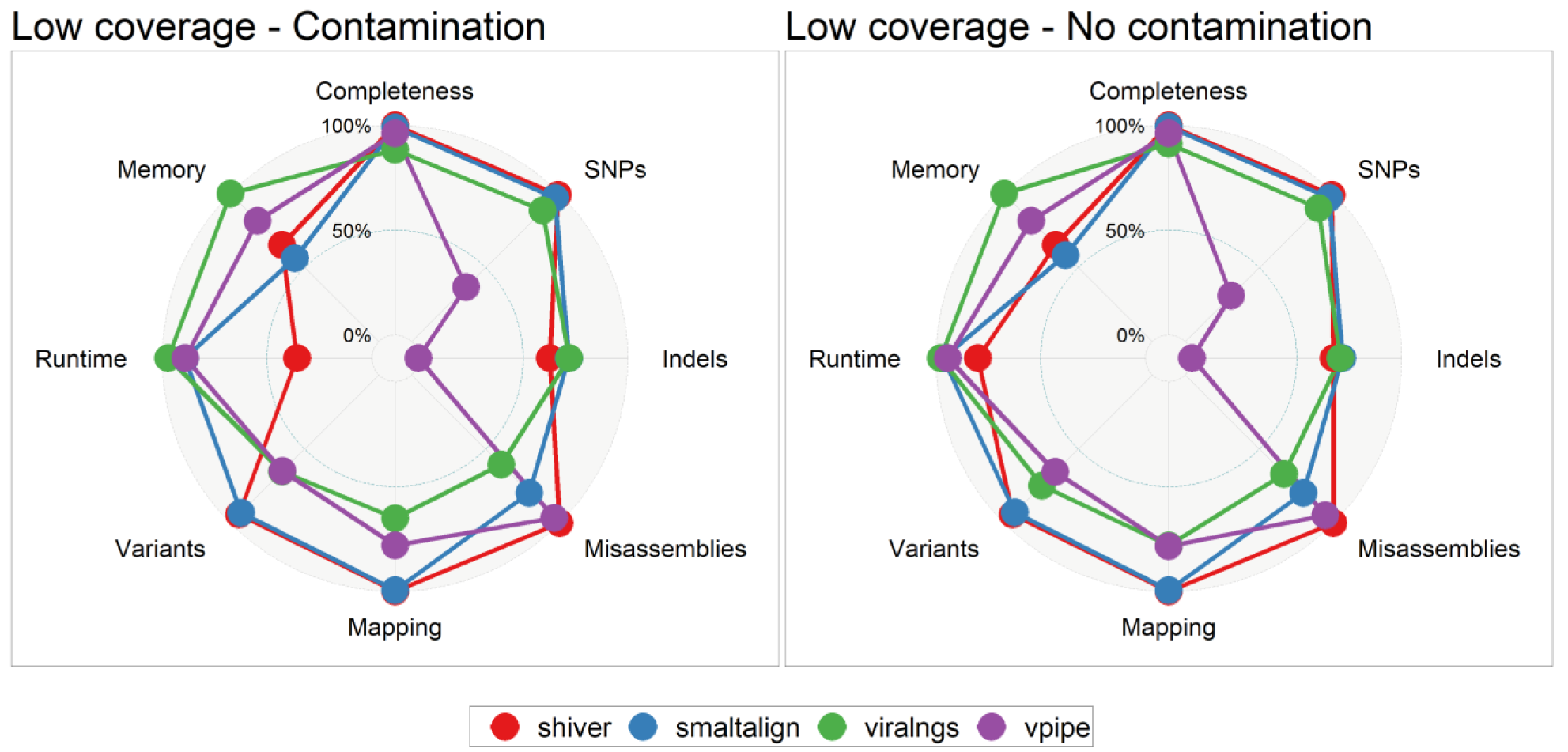
Multidimensional performance of genome assemblers in low coverage scenarios. Benchmarking metrics are compared for A) the low coverage – contamination and B) for the low coverage – no contamination scenarios of the SIM dataset. For an explanation on the scaling of the different axes see the figure caption of Figure S6.

## Biographical Notes

Levente Zsichla is finishing his Master’s degree in Biology with a specialization in Bioinformatics, and is a research assistant at the National Laboratory for Health Security, at Eötvös Loránd University, Budapest, Hungary.

Marius Zeeb is a PhD student at University Hospital Zurich, Switzerland, focusing on the epidemiology of HIV-1 and tuberculosis.

Dávid Fazekas is a data scientist (and currently also security engineer and DevOps specialist) with a research background in bioinformatics.

Éva Áy is a molecular biologist at the National Center for Public Health and Pharmacy, Budapest, Hungary and a research associate at the National Laboratory for Health Security, at Eötvös Loránd University, Budapest, Hungary. She currently leads the National Reference Laboratory for Retroviruses.

Dalma Müller received her MSc degree in Biology with a specialization in Molecular Genetics, Cell- and Developmental Biology from Eötvös Loránd University, Budapest, Hungary. She is currently a PhD student at Semmelweis University, Budapest, Hungary, and a research associate at the National Laboratory for Health Security, Eötvös Loránd University, Budapest.

Karin J. Metzner is an adjunct professor of molecular virology at the University of Zurich, Switzerland, focusing on the genomic organization, diversity, and evolution of HIV-1, with implications for pathogenesis, drug resistance, latency, and transmission.

Roger Kouyos is an associate professor at the University Hospital Zurich, Switzerland, studying infectious diseases by integrating approaches from epidemiology, bioinformatics, and evolution.

Viktor Müller is an associate professor and PI of the National Laboratory for Health Security at Eötvös Loránd University, Budapest, Hungary. His research interests encompass the modeling, data analysis and evolution of infectious diseases.

